# Analysis of a genetic region affecting mouse body weight

**DOI:** 10.1101/2022.08.24.502873

**Authors:** Connie L.K. Leung, Subashini Karunakaran, Michael G. Atser, Leyla Innala, Xiaoke Hu, Victor Viau, James D. Johnson, Susanne M. Clee

**Author notes:** Correspondence to: James D. Johnson, Ph.D. Life Sciences Institute Diabetes Research Group. The University of British Columbia | Point Grey Campus | Musqueam Traditional Territory 5358 – 2350 Health Sciences Mall | Vancouver, BC, Canada V6T 1Z3 | @JimJohnsonSci. Deceased (Susanne M. Clee). **Dedication:** We dedicate this work to the memory of Dr. Susie Clee, who is dearly missed.

## Abstract

Genetic factors affect an individual’s risk of developing obesity, but in most cases each genetic variant has a small effect. Discovery of genes that regulate obesity may provide clues about its underlying biological processes and point to new ways the disease can be treated. Pre-clinical animal models facilitate genetic discovery in obesity because environmental factors can be better controlled compared to the human population. We studied inbred mouse strains to identify novel genes affecting obesity and glucose metabolism. BTBR T+ Itpr3tf/J (BTBR) mice are fatter and more glucose intolerant than C57BL/6J (B6) mice. Prior genetic studies of these strains identified an obesity locus on chromosome 2. Using congenic mice, we found that obesity was affected by a ~316 kb region, with only two known genes, pyruvate dehydrogenase kinase 1 (Pdk1) and integrin alpha 6 (Itga6). Both genes had mutations affecting their amino acid sequence and reducing mRNA levels. Both genes have known functions that could modulate obesity, lipid metabolism, insulin secretion and/or glucose homeostasis. We hypothesized that genetic variation in or near *Pdk1* or *Itga6* causing reduced *Pdk1* and *Itga6* expression would promote obesity and impaired glucose tolerance. We used knockout mice lacking *Pdk1* or *Itga6* fed an obesigenic diet to test this hypothesis. Under the conditions we studied, we were unable to detect an individual contribution of either *Pdk1* or *Itga6* to body weight. However, we identified a previously unknown role for *Pdk1* in cardiac lipid metabolism providing the basis for future investigations in that area.

## Introduction

Obesity and type 2 diabetes are complex diseases that share common risk factors. These metabolic diseases share inter-related traits such as increased BMI, adiposity, insulin resistance, and mitochondrial dysfunction. Discovery of genetic factors affecting these traits in humans has been productive but challenging, due to the relatively small effects of many genetic variants, and the effects of inter-individual variation in environmental and lifestyle factors that can have significant impact on the phenotypic expression of these genetic changes. Genome wide association studies (GWAS), and advances in sequencing technology have allowed further studies to be conducted to find polygenic obesity genes that affect obesity susceptibility in the general population. However, the known common variants affecting the general population have modest effects on body weight. For example, GWAS data from a cohort of 694,649 samples found 346 loci associated with BMI had relatively small individual effect sizes, and the combined loci only accounted for ~3.9% of the variation in BMI (Pulit et al. 2019). In addition to genetic factors, lifestyle factors such as stress (Koch, Sepa, and Ludvigsson 2008; Tomiyama 2019) availability to and access to healthy food, or socioeconomic status (Volaco, Cavalcanti, and Précoma 2018) have been associated with obesity. This complexity confounds the discovery of obesity genes in humans.

As an alternative approach to overcome some of these challenges, we have capitalized on the natural genetic differences that occur between inbred mouse strains, and the strong evolutionary conservation of obesity mechanisms across species. The BTBR T+ *Itpr3^tf^*/J (BTBR) mouse strain has a high propensity for obesity. Prior studies have shown that this strain harbours alleles that promote metabolic disease compared to the C57BL/6J (B6) strain (Clee, Nadler, and Attie 2005; Ranheim et al. 1997; Stoehr et al. 2004a; Stoehr et al. 2000). Genetic studies identified several quantitative trait loci (QTLs) that affect metabolic disease traits in these strains (Stoehr et al. 2004a; Stoehr et al. 2000), and genes affecting these traits have been identified (Bhatnagar et al. 2011; Clee et al. 2006). Additional insights about factors affecting metabolic disease in these strains have come from multiple ‘omic studies and their integration with genetics (Davis et al. 2010; Ferrara et al. 2008; Keller et al. 2008; Lan et al. 2003; Nadler et al. 2000; Tian et al. 2015; Tu et al. 2012; Zhao et al. 2009; Dong et al. 2021).

One quantitative trait locus (QTL) identified using these strains is *Modifier of obese 1* (*Moo1*)(Stoehr et al. 2004a). At this locus, BTBR alleles were associated with increased body weight. We have previously localized this QTL to a ~6 Mb region of chromosome 2, and have shown that it is associated with multiple metabolic traits (Karunakaran et al. 2013; Stoehr et al. 2004a). Here, we report that *Moo1* is comprised of at least 2 QTLs, localized the major effect to a 316 kb region of chromosome 2, described the environmental modification of this locus by stress, as well as insights into lipid metabolism when PDK1 (pyruvate dehydrogenase kinase 1) levels are reduced.

## Methods

### Animals

Animals were housed in ventilated microisolator cages in environmentally controlled facilities with 12 hr light:dark cycles, unless otherwise noted. All mice were generated from crosses between heterozygotes (HETs) such that littermates were used as controls. Congenic mice with B6 regions of chromosome 2 replacing those of the BTBR background strain were generated from an in-house breeding colony. Subcongenic strains were created from recombinants of the Moo1C strain previously described (Karunakaran et al. 2013). A heterozygous breeder pair of *Pdk1*^tm1.1(KOMP)Vlcg^ knockout (KO) mice were obtained from the Knockout Mouse Project (KOMP), then bred in house. These mice were originally on the C57BL/6N background but were backcrossed to C57BL/6J as part of colony expansion and maintenance. Because prior studies showed no effect of this locus in females, experiments used exclusively male animals.

Over the course of these studies, mice were housed in multiple facilities and rooms within these facilities, Center for Disease Modelling (CDM) room 1, room 2 and room 3 and Wesbrook Mouse Facility. The majority of the studies were completed within the CDM at the University of British Columbia (UBC). Analysis of the subcongenic strains was performed concurrently with mice housed in room 1 in CDM. However, CDM underwent reorganization, and the mice were moved to CDM room 2. The stress study was conducted in CDM room 2. CDM then underwent construction and thus when CDM reopened, a repeat analysis of Moo1V strain concurrent with ITGA6 KO and repeat analysis PDK1 KO in CDM room 3 was conducted.

At weaning, all experimental animals were placed on a diet high in fat with sucrose (D12492i, Research Diets). Mice were weighed weekly, at a standardized time of day. At the specified times, mice were euthanized by overdose of isoflurane anesthesia in accordance with the Animal Care guidelines. At this time a cardiac blood sample was withdrawn, and tissues were rapidly harvested, weighed and flash frozen. Blood samples were kept on ice then plasma was separated by centrifugation at 4°C at 10,000 rpm for 8 minutes. Tissue and plasma samples were stored at −80°C until analysis. All procedures were approved by the University of British Columbia Animal Care and Use Committee and were performed in accordance with Canadian Council on Animal Care Guidelines (Protocol # A19-0267).

### Genotyping

DNA was extracted from ear notches using a commercially available kit (Puregene, Qiagen). Congenic mice were genotyped at the first and last markers shown to be B6 within the congenic insert. Recombinants were confirmed, then additional markers genotyped to fine-map the recombination. Genotyping of SNP markers was performed using high resolution melt curve analysis (Qiagen kit) using the Rotor-gene Q thermocycler (Qiagen). Microsatellite markers were genotyped by PCR amplification and resolution on polyacrylamide gels, as described(Skarnes et al. 2011). PDK1 KO mice were genotyped by PCR amplification of of a 195 bp band for the WT allele and a 639 bp band for the KO allele as recommended by KOMP (Skarnes et al. 2011). Genotyping of *Itga6* mice was conducted with primers flanking the region of the loxP sites to amplify a fragment of 120 bp in WT animals and, 150 bp when the loxP site is present. Genotyping for the presence of cre recombinase was performed using a set of primers that amplify across the deleted region were used to amplify a fragment of 600 bp, as described (Georges-Labouesse et al. 1998). Primer sequences are provided in **Supplemental Table 1**. All genotyping reactions contained DNA samples known to be of each genotype as controls, along with a no template control.

### Gene expression analysis

Tissue samples were processed, and RNA was extracted using E.Z.N.A total RNA extraction kit (Omega Bio-tek, Norcross, GA), after which cDNA was generated using the RevertAid First Strand cDNA Kit (ThermoFisher Scientific, Cat #K1631, Waltham, Massachusetts, United States). Gene expression was measured using real time qPCR with SYBR green. β-actin (*Actb*) was used as a reference gene as it more consistently amplified in the same manner across different experimental groups and tissues compared to Glyceraldehyde 3-phosphate dehydrogenase (*Gapdh*) or cyclophilin (*Ppib*). A single product was verified using melt curve analysis. Cycle threshold value (Ct) was subtracted from the control gene Ct value, to give the Delta Ct value (dCt). Then delta delta Ct values (ddCt) for each mouse were calculated by subtracting each dCt value from the average dCt of the control group. No reverse transcriptase (no RT) and no template control (water) were used as negative controls and were determined for each gene and tissue assessed.

### Body weight and blood sampling

Mice were weighed weekly at a standardized time of day from weaning at 3 weeks until 10 weeks of age. To determine changes in body composition, dual-energy X-ray absorptiometry (DEXA) analysis was performed using a PIXImus Mouse Densitometer (Inside Outside Sales, Madison, WI). Body weight, body length, and cardiac blood samples were also obtained at sacrifice.

### Stress study

To directly assess the interaction between stress and genotype on obesity, we used chronic restraint stress, a well-established model of stress in rodents(Ball et al. 2017). At 10 weeks of age, the mice were singly housed for the duration of the study. At 11 weeks of age, restraint stress was induced in half the mice by placing the mice in a plexiglass restrainer for 90 minutes from 9 am to 10:30 am daily, for 7 days. The plexiglass restrainer confined the mice into a small space, with no room to move, but it did not specifically immobilize them. The control mice were singly housed during the week of stress but were not placed in the restrainers; however, all the other hormone and metabolic measurements were performed concurrently with the stress group. Age matched littermates were used in both the control and stressed groups. To assess food intake during the experiment, mice were given a pre-weighed amount of food at the start of the study. The weight of the food remaining at the end of the study was determined again to calculate a total food intake during the 1 week of restraint stress. There were no visible crumbs of food in the cages when the food weight was measured. Corticosterone levels were measured in plasma from the cardiac bleeds of the mice using a Corticosterone Double Antibody RIA kit (MP Biomedicals, Solon, OH) as previously described (Goel, Innala, and Viau 2014). Thirty-six hours after the last stress induction, the mice were euthanized and a standard set of tissue samples for all of our mice were collected, including brain, heart, stomach, intestines, spleen, fat deposits (epididymal, renal, mesenteric, brown adipose tissue (BAT)), soleus muscle, and testes. These tissues were weighed and then flash frozen on dry ice as soon as they were collected. Immediately after euthanasia, prior to tissue collection, cardiac blood samples were collected and placed into eppendorf tubes containing 6 μL of 25mM EDTA and kept on ice until plasma separation. All plasma samples were separated by centrifugation at 4°C 10,000 rpm for 8 minutes, and all tissue and plasma samples were stored at −80°C until analysis.

### Glucose and Triglycerides

Metabolite levels were measured in plasma from the cardiac blood samples collected from mice at sacrifice. The mice were fasted for 4 hours before sacrifice. Glucose levels were measured with a colorimetric assay as described above (Autokit Glucose Cat# 997-03001, Wako Diagnostics, Richmond, VA, USA). Insulin was measured by ELISA (Mouse Ultrasensitive Insulin ELISA, Cat#80-INSMSU-E01, Alpco, Salem, NH, USA). Plasma β-hydroxybutyrate levels (Beta Hydroxybutyrate Assay kit, Cat# ab83390, Abcam, Cambridge, MA, USA), cholesterol (Cholesterol-SL Assay, Cat# 234-60 Sekisui Diagnostics, PEI, Canada), triglyceride and glycerol (Triglyceride-SL Assay, Cat# 236-60, Sekisui Diagnostics, PEI, Canada) and pyruvate (Pyruvate Assay Kit, Cat#MAK071, Sigma-Aldrich, Oakville, ON, Canada) were measured using colorimetric assays according to the manufacturers’ directions. The products of the colorimetric assay were measured spectrophotometrically at 505 nm for glucose and cholesterol, at 520nm for triglycerides, at 450 nm for insulin and β-hydroxybutyrate, and at 570 nm for pyruvate using an Infinite M1000 microplate reader (Tecan, Durham, NC, USA).

Tissue cholesterol and triglyceride levels were measured in liver and heart samples (Briaud et al., 2001). Tissues (approximately 100 mg) were homogenized using a tissue homogenizer in 3 mL of chloroform methanol (2:1), and extracted using 1.5 mL of ice-cold water and a second time with 750 μL of ice cold water. A third of the measured organic layer from the liver extraction was dried with 30 μL of Thesit (hydroxypolyethoxydodecane) neat (Sigma-Aldrich, Oakville, ON, Canada) under nitrogen gas. The volume used for dehydration was measured precisely after the aliquot, as to not disturb the organic layer. For the heart samples, the entire organic layer was dried. Standards with cholesterol (Wako Diagnostics, Richmond, VA, USA) and triolein (1:250 uL) (Sigma-Aldrich, Oakville, ON, Canada) were dried under nitrogen gas. Once dried, the samples were mixed with 300 μL of water and incubated at 37°C with vigorous vortexing twice partway through the 30-minute incubation. Samples were stored at 4°C until analysis. Before the analysis, samples were brought up to room temperature and diluted with 10% Thesit as needed, as determined by first testing undiluted samples then repeating the assay on samples that were not on the standard curve (Huynh et al. 2010).

Tissue cholesterol was measured using a colormetric assay (Cholesterol E, Cat #439-17501, Wako Diagnostics, Richmond, VA, USA). The cholesterol reagents were reconstituted according to the manufacturer’s instructions. In a flat bottom 96 well plate (Microtest plate 96 well, Cat#82.158, Sarstedt, Nümbrecht, Germany), 20 μL of extracted lipids and standards were added, then 200 μL of cholesterol E reagent (Cholesterol E, Cat #439-17501, Wako Diagnostics, Richmond, VA, USA) was added and incubated at room temperature for 5 minutes. The products of the colorimetric assay were measured spectrophotometrically at 590 nm using an Infinite M1000 microplate reader (Tecan, Durham, NC, USA). The values were expressed per amount of starting protein and accounting for the dried volume.

Tissue triglyceride levels were measured using an enzymatic triglyceride assay (Glycerol Reagent, Cat# F6428, Triglyceride Reagent, Cat#T2449, Sigma-Aldrich, Oakville, ON, Canada). In a flat bottom 96 well plate, (as described above), 5 μL of extracted lipids and standards and 200 μL Glycerol Reagent (Glycerol Reagent, Cat# F6428, Sigma-Aldrich, Oakville, ON, Canada) were added and first measured to obtain a reading for glycerol at 540nm. Then 50 μL of Triglyceride Reagent (Triglyceride Reagent, Cat#T2449, Sigma-Aldrich, Oakville, ON, Canada), which contains lipase to release glycerol from the fatty acids, was added to each of the samples. The absorbance values of the plate were measured at 540 nm using an Infinite M1000 microplate reader (Tecan, Durham, NC, USA). The samples were incubated for 5 minutes at 37°C before each absorbance reading. The last absorbance read was then subtracted from the initial plate absorbance to obtain the final absorbance due to glycerol from triglycerides.

To examine triglyceride synthesis and secretion in Pdk1 KO mice, a standard lipogenesis test (triglyceride synthesis and secretion) was conducted using poloxamer 407 (Millar et al. 2005) at 12 weeks of age. Poloxamer 407 is a chemical that blocks triglyceride breakdown and clearance from the blood by inhibiting lipoprotein lipase (Millar et al. 2005). Thus, during fasting, increasing plasma triglyceride concentrations result from increased triglyceride synthesis and secretion. The mice were fasted for 4 hours prior to testing. Blood was collected at time 0, and 30, 60, 120, 240 and 360 minutes post treatment of injection with poloxamer 407 (1g/kg intraperitoneal). Saphenous blood was collected into tubes containing 6 μL of 25mM EDTA and kept on ice until plasma separation. All plasma samples were separated by centrifugation at 4°C 10,000 rpm for 8 minutes. Plasma was stored at −80°C until further analysis.

To assess lipid clearance from circulation in Pdk1 KO mice, an oral lipid tolerance test was performed at 14 weeks of age. Mice were fasted overnight (5pm-9am). At the time food was removed the mice were placed in clean cages with water prior to the lipid tolerance tests and kept in these cages throughout the testing to ensure that the triglycerides in the plasma were from the oral gavage instead of from food crumbs. For the lipid tolerance, saphenous blood was collected at time 0, 30, 60, 120, 240 and 360 minutes post oral gavage of olive oil (300 μL) (Excoffon et al. 1997). Saphenous blood was collected into tubes containing 6 μL of 25mM EDTA and kept on ice until plasma separation. All plasma samples were separated by centrifugation at 4°C 10,000 rpm for 8 minutes and stored at −80°C until further analysis.

### Statistical analyses

Statistical analyses, including 2-way ANOVA and Student’s T-test, were performed using GraphPad Prism Software (Version 6.0, La Jolla, CA, USA). A *p* value of less than 0.05 was considered significant.

## Results

### Localization of Moo1

Previous studies in the Dr. Alan Attie’s laboratory at the University of Wisconsin had mapped a region that causes an increase in body weight in BTBR mice compared to B6 mice on the ob/ob background, despite the extreme hyperplasia of the mice without functional leptin, to chromosome 2. Fine mapping was done using congenic mice to localize a region on chromosome 2 that causes an increase in body weight when BTBR alleles are inherited (Stoehr et al. 2004b). The chromosome 2 region found in the ob/ob mouse background was also found to affect HFD-induced obesity. A recent independent genetic cross also showed a significant linkage peak in this region (Suppl. Fig. 1) (Tian et al. 2015; Tu et al. 2012) We have previously described the localization of *Moo1* to a ~6 Mb region of chromosome 2 contained within a congenic strain, Moo1C (Karunakaran et al. 2013). To further genetically localize *Moo1*, we created a panel of subcongenic strains from recombinants of the Moo1C strain (Fig. 1).

**Figure 1.**
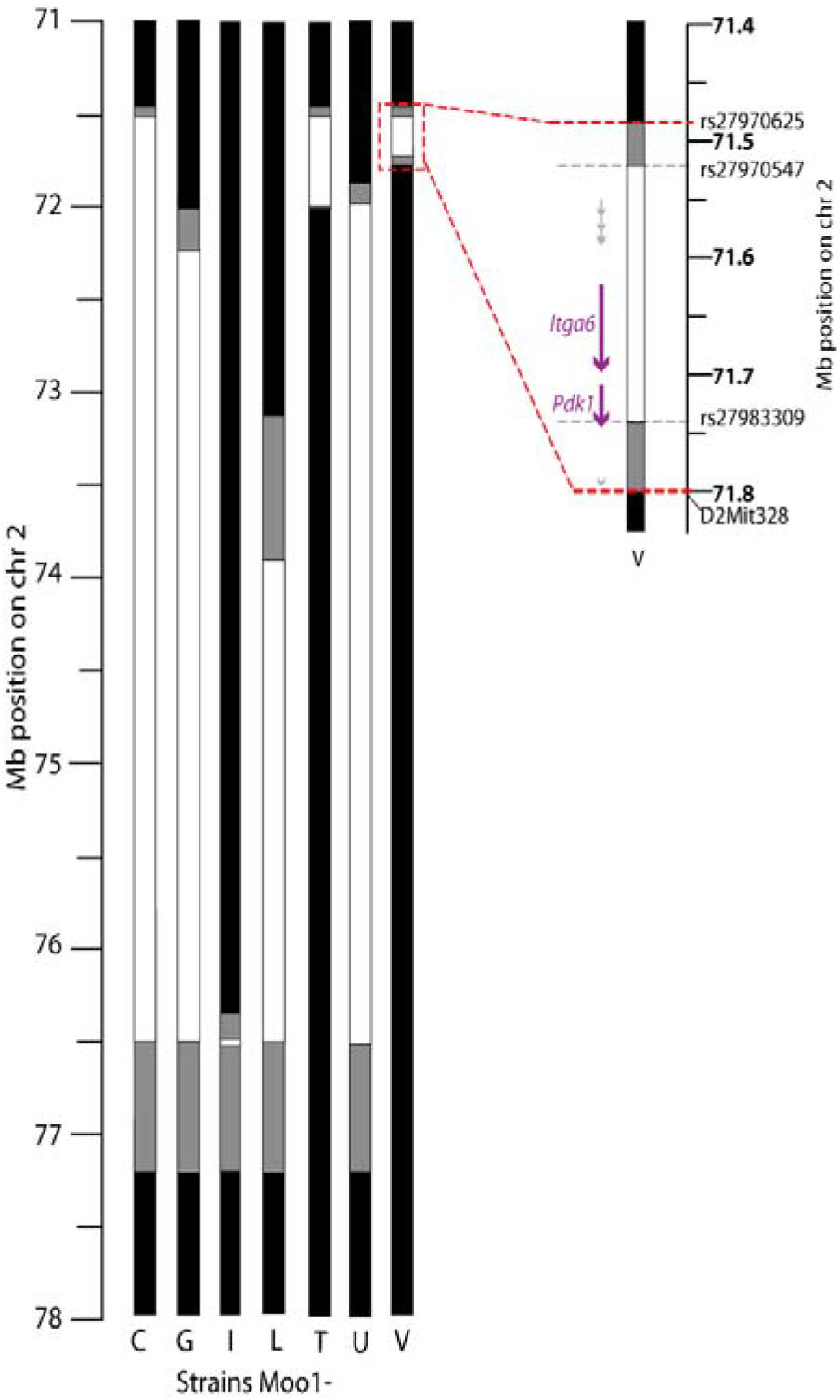
Moo1 subcongenic strains. Subcongenic strains with B6 congenic inserts in a BTBR genomic background, drawn to scale. Positions on chromosome 2 are from the mm9 genome assembly. The markers defining the boundaries of the Moo1V strain are shown on the zoomed version. White boxes represent the B6 congenic insert, grey boxes are undetermined genotype between the last markers tested as B6 and the first marker tested as BTBR, black = BTBR region as throughout the rest of the genome. On the zoomed version, the location of the known genes (*Itga6*, *Pdk1*; purple) and predicted genes (from GENCODE VM20; grey). The upper grey line includes 5 predicted genes: Platr26, Gm13647, Gm13663, Gm17250, Gm13662; the grey arrow at the bottom of the region is Gm13746. None of these predicted genes are shown to have orthologues in other species, including rat which is very closely related to mouse.

Before these mice were moved to the Clee laboratory at UBC, the congenic mice were backcrossed to remove the ob/ob allele background. We assessed body weights weekly in mice from each strain, as in our prior studies in the genetic screens (Fig. 2) (Karunakaran et al. 2013). These studies found that at least 2 independent loci within the Moo1C region contributed to the QTL, even without the ob/ob allele background. A major locus localized to a 316 kb region in the Moo1V strain, and has been termed *Moo1a*. A second locus having a smaller effect was localized telomeric to this region in the non-overlapping Moo1G strain, and has been termed *Moo1b*. For the studies described in this paper, we focused on Moo1V strain (*Moo1a*).

**Figure 2.**
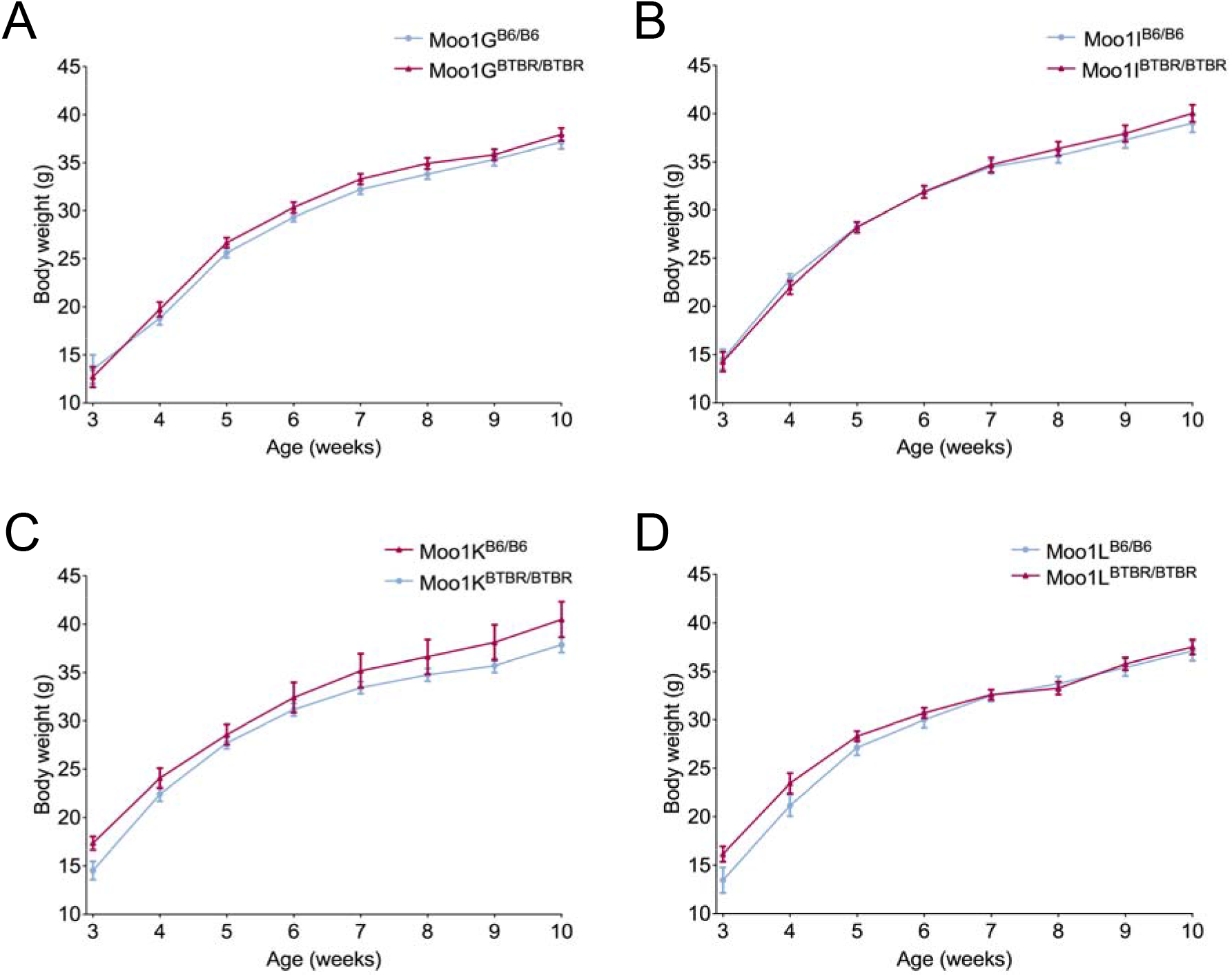
Body weight analysis of Moo1 subcongenic strains. Male mice were fed the same HFD, housed in CDM room 1 with the Moo1C and Moo1V mice, and weighed weekly. Neither (A) Moo1G strain mice, (B) Moo1i strain mice, (C) Moo1K strain mice, or (D) Moo1L strain mice had significant differences in body weights between genotypes. Data are mean ± SEM analyzed by two-way ANOVA. ns denotes not significant.

To identify genetic differences between the strains, we designed primers flanking each exon and spanning ~2 kb of promoter for *Itga6* and *Pdk1*, and performed Sanger sequencing. Numerous differences were identified (Table 1). Notably, within the coding region of *Itga6* was a nonsynonymous change that results in a leucine in B6 mice and a valine in BTBR (now rs13464795). Notably, valine is present in many other species, suggesting it may be the ancestral allele (Suppl Fig 2). Within the coding region of *Pdk1* was a 6 bp insertion/deletion (indel) that results in the presence of an alanine and serine at amino acids 27 and 28 in B6 mice, but their absence in the BTBR strain (now rs222188523). Interestingly, the N-terminal region of PDK1 varies in length across species (Suppl Fig 3), so the functional importance of this change is unclear, however in the mouse strains with sequence data available, these two amino acids appear only in B6 (Suppl Fig 3&4). In both genes, we also identified many variants in non-coding regions, synonymous variants, and variants in the untranslated regions (Table 1). Since this work was performed, many mouse strains in addition to B6 have been sequenced, including BTBR. These data, retrieved from the Mouse Phenome Database, list a total of 3416 variants (SNPs and indels) differing between the B6 and BTBR strains in the 316 kb region of the Moo1V strain. Because we identified many non-coding variants around *Itga6* and *Pdk1*, we examined whether these may affect their expression in multiple tissues. These studies showed that Moo1^BTBR/BTBR^ mice, have a ~50% reduction in gene expression of *Pdk1* and *Itga6* in most tissues compared to Moo1^B6/B6^ mice (Fig. 3). These data are consistent with a recent genetic analysis of these strains that identified cis-eQTLs (Tian et al. 2015; Tu et al. 2012) for both *Pdk1* and *Itga6* (Suppl Fig. 1). Thus, we considered both *Pdk1* and *Itga6* as positional candidate genes.

**Table 1.** Sequence differences between B6 and BTBR in *Pdk1 and Itga6*.

| Gene | 2 kb promoter | in mRNA |  |  |  |
| --- | --- | --- | --- | --- | --- |
|  |  | non-synonymous | synonymous | untranslated region | intronic |
| *Pdk1* | 8 | delAla, Ser 27,28 | Leu96<br>Asp99<br>Asp426 | 83 | 23 |
| *Itga6* | 11 | Leu975Val | Arg91<br>Val119<br>Leu157<br>Thr243<br>Val290<br>Ala299<br>Ile333<br>Leu522<br>Leu952 | 1 | 39 |

Data showing the number of changes in BTBR mice in each gene, by location, obtained by Sanger sequencing performed by the Clee Lab. The first ~2 kb of the promoter was sequenced, and only short segments of each intron immediately flanking the exons were sequenced. There is a 6 nucleotide deletion resulting in the deletion of an alanine and serine in PDK1. A mutation that results in the change of a leucine to valine in ITGA6 was also found. The number of variants in the promoter, introns, and mRNA (that were non-synonymous, synonymous or in the untranslated region) are shown. Data obtained subsequently from the Jax Mouse Phenome Database website (www.jax.org/phenome, Sanger4 project) from large scale genotyping confirmed these variants and identified 2922 sequence variants between B6 and BTBR in the entire 316 kb region.

**Figure 3.**
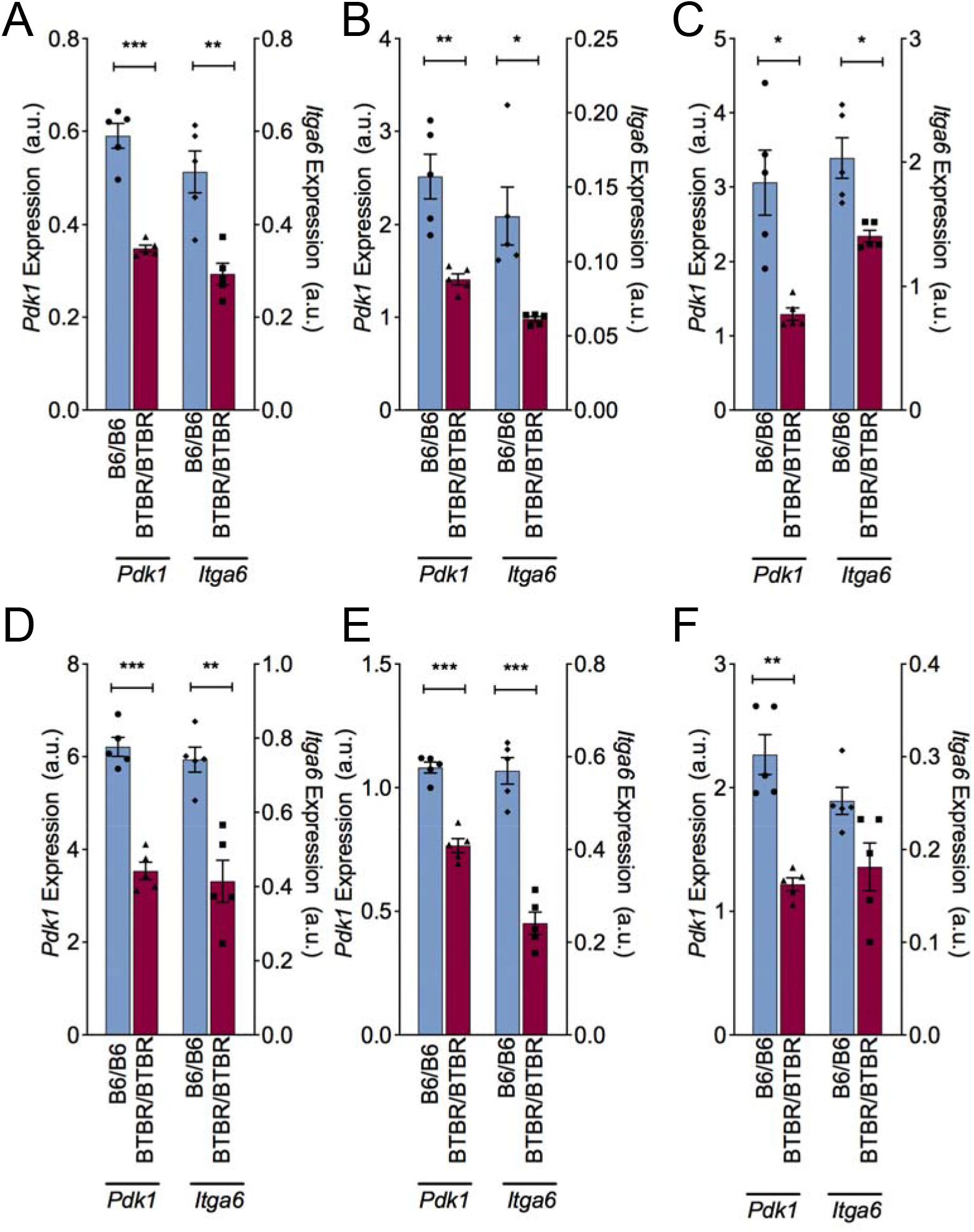
Gene expression of Pdk1 and Itga6 is reduced in BTBR and B6 parental strains. Gene expression in (A) Hypothalamus (B) Liver (C) Adipose tissue from epididymal fat pad (D) Soleus muscle (E) Gastrocnemius muscle (F) Islets from 4 week old B6 (red, n= 5) or BTBR (blue, n= 5) mice fed chow at UW. Data are raw microarray expression values, obtained from diabetes.wisc.edu, and are shown as the mean± SEM. Statistics were performed by student’s T test (*p<0.05, **p<0.01, ***p<0.001).

### Environmental and stress modulation of Moo1

During our studies with the Moo1C strain, the mice were relocated to a new university (UBC) and multiple housing locations (see above in the Methods section). The effect of *Moo1* on body weight in these strains varied between locations, even becoming undetectable in one location (Fig. 4A-C). Subsequently, similar environmental effects were observed for the Moo1V strain housed in different rooms in the same facility (Suppl. Fig. 5). Collectively, these observations indicate that gene-environment interactions likely play a strong role in the penetrance of this phenotype.

**Figure 4.**
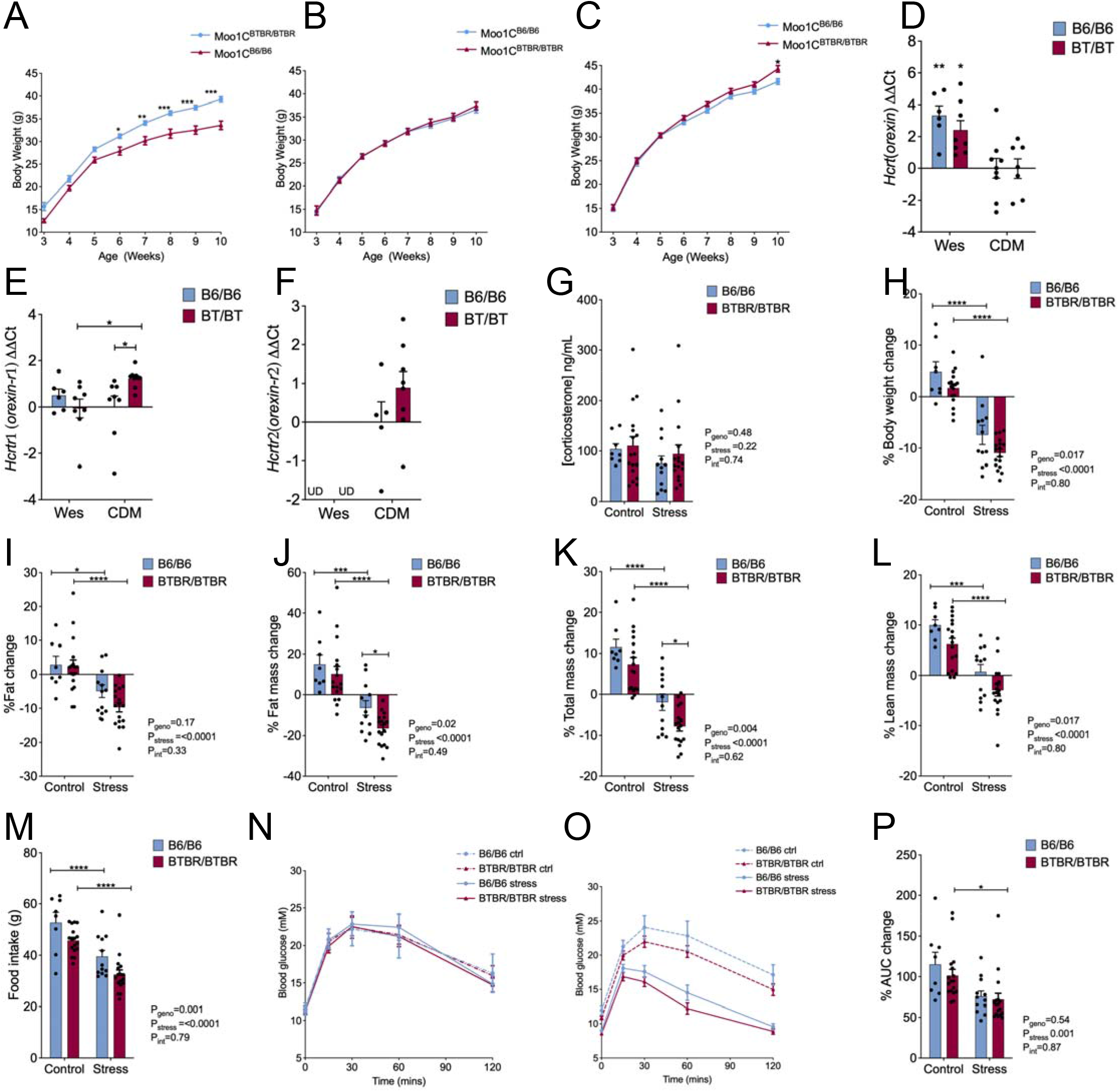
Environmental and stress modulation of the body weight phenotype. Mice were moved from the University of Wisconsin-Madison (UW) to the University of British Columbia (UBC-Wesb) and into a second facility at UBC (UBC-CDM). (A) Moo1C growth curves in UW, (B) Wesbrook, and (C) CDM facilities. (D) Orexin (*Hcrt*), (E) Orexin receptor 1 (*Hcrtr1*) and (F) Orexin receptor 2 (*Hcrtr2*) expression levels in the brain were assessed in Moo1C congenic mice from the Wesbrook (Wesb) facility compared to the CDM facility. Data are ΔΔCt (ddCt) values relative to Moo1C^B6/B6^ mice in CDM. n=6-9. (G) Corticosterone levels in cardiac blood samples of Moo1V strain mice at 12 weeks of age did not differ. Mice corticosterone levels were not significantly different in all four groups. Mice were housed in CDM room 2 and were fed a HFD. (H) Body weight was assessed in 12 weeks mice 24 hours before and 24 hours after the last stress treatments. Moo1V^B6/B6^(B6/B6) control n=8, stress n=12; Moo1V^BTBR/BTBR^ (BTBR/BTBR) stress and control group n=17 per group. Body composition was assessed by DEXA before and after the one week experiment. Percent change in (I) percent body fat, (J) fat mass, (K) total mass, (L) lean mass. Moo1V^B6/B6^ control n=8, Moo1V^B6/B6^ stress n=12, Moo1V^BTBR/BTBR^ stress n=17, and Moo1V^BTBR/BTBR^ control group n=17. (M) Food intake was assessed in each group during the week of stress, day 7 food weight was subtracted from that given on day 1 of the experiment. Moo1V^B6/B6^ control n=8, Moo1V^B6/B6^ stress n=12, Moo1V^BTBR/BTBR^ stress n=17 and Moo1V^BTBR/BTBR^ control group n=17. GTTs were conducted both 24 hours before and after the 1 week of stress treatment and at the same time in non-stressed controls. Glucose tolerance in mice (N) before and (O) after the stress treatment were measured and (P) AUC was calculated as area under the curve to t=0. Moo1V^B6/B6^ (B6/B6) control n=8, Moo1V^B6/B6^ stress n=12, Moo1V^BTBR/BTBR^ (BTBR/BTBR) stress n=17 and Moo1V^BTBR/BTBR^ control group n=17. Data are shown as mean percent change in each group ± SEM and were analyzed by two-way ANOVA with stress treatment and genotype as the factors (shown below each graph, along with the p-value for their interaction). Bonferroni pairwise comparisons between groups that were significantly different are shown (*p<0.05, ***p<0.001, ****p<0.0001) or ns for not significant.

One notable difference between the housing locations was the noise and potential disturbances to the mice due to adjacent construction. We assessed the expression of genes involved in pathways by which stress is known to affect food intake in brains from Moo1C mice housed at the two UBC facilities, CDM and Wesbrook. Orexin expression levels were 4- to 8-fold higher in mice housed in the Wesbrook facility compared to the mice housed in the CDM, suggesting that differences between the facilities affected this pathway (Fig 4D-F).

To directly test whether stress modifies the effect of *Moo1* on body weight, we conducted controlled stress experiments to identify stress effects on body weight and composition, food intake and glucose tolerance. To determine the degree to which mice were stressed due to their placement in the restrainers, cortisol levels were measured at the end of the study. Corticosterone levels were the same in both the control and the stress group, suggesting that the mice did not respond differently to stress (Fig 4G). The mean levels in all groups were elevated compared to what is found in normal, non-stressed mice, typically <46 ng/mL and stressed levels are typically >100 ng/mL (Park et al. 2017; Valentine, Williams, and Maurer 2012; Flint and Tinkle 2001; Shanks et al. 1990). To measure food intake, the mice were singly housed throughout the study, which could increase stress in all groups. In addition, the corticosterone levels were measured in blood collected 36 hours after the last stress induction. This collection would be after the expected peak corticosterone levels(Goel et al. 2022), which may explain the smaller differences in the control compared to the stress groups, resulting in no differences between the groups. Body weight increased in the non-stressed control group during this one-week period. In contrast, their littermates which were subjected to stress lost weight. These effects were not significantly different between genotypes (Fig 4H). We also assessed the effects of stress on body composition. Stress significantly reduced fat mass (Fig 4J) and lean mass (Fig 4L) in both genotypes. The effect on fat mass was proportionally greater, resulting in significant decreases in percent body fat (Fig 4I). Notably, both total mass and fat mass were reduced significantly more in response to stress in the Moo1V^BTBR/BTBR^ genotype compared to Moo1V^B6/B6^ mice (Fig 4J &K). In the non-stressed control mice, the change in these parameters did not differ between genotypes. This suggests that stress affects each genotype differently. Food intake was significantly reduced in Moo1V^BTBR/BTBR^ mice compared to Moo1V^B6/B6^ mice (Fig 4L). The difference between genotypes was similar in both control and stress groups. As we expected with a decrease in adiposity in the stressed mice, we found that under stress, the Moo1V mice had improved glucose tolerance (Fig 4N-P).

Our observations that the *Moo1* phenotype is modulated by housing environment, and body weight analysis was performed in CDM room 2 before we obtained the Pdk1 and Itga6 mice, we repeated the Moo1-V strain body weight analysis in the newly renovated facility, CDM room 3. This analysis was performed in the same location and simultaneously with Pdk1 and Itga6 mice. In this housing location, although the effects of *Moo1a* on body weight were blunted, body fat was still reduced in mice with the Moo1V^B6/B6^ mice (Suppl Fig. 5).

### Effects of reduced Pdk1 on body weight

*Pdk1* encodes pyruvate dehydrogenase kinase 1. There are four kinases in this family that phosphorylate and inhibit pyruvate dehydrogenase (PDH), blocking the conversion of pyruvate to acetyl-CoA for entry into the Krebs Cycle for glucose metabolism (Korotchkina and Patel 2001a; Korotchkina and Patel 2001b). Reduced PDK1 is expected to increase acetyl-CoA formation. Acetyl-CoA is a substrate for lipogenesis and its activation could thereby promote fat storage. Thus, *Pdk1* is both a positional and functional candidate gene, with a reduction in PDK1 activity expected to promote obesity. We tested this hypothesis using global *Pdk1* KO mice.

We first analyzed the role of *Pdk1* KO in body weight and measured weekly body weights to 10 weeks of age. Growth curves from *Pdk1* KO mice were not significantly different from those of their HET or wildtype (WT) littermate controls (Fig. 5A). A lack of difference in total body weight does not necessarily indicate that there are no changes in adiposity. We measured adiposity as well. We did not detect a significant difference in body composition measured by DEXA in the KO mice compared to littermate HET and WT controls (Fig. 5A). We also measured body length to identify if there were differences to body length, but found no significant differences (Fig. 5B). However, we found increases in epididymal and mesenteric fat in the Moo1V^BTBR/BTBR^ mice compared to Moo1V^B6/B6^ mice (Fig. 5E&F) when differences in total body fat were not apparent. Thus, we also weighed fat pads at sacrifice. Fat pad weights were not significantly different between genotypes except for mesenteric fat, which was found to weigh less in KO mice compared to HET (Fig. 5F). This contrasts with our expectation from the congenic mice, where the genotype with reduced *Pdk1* expression had increased adiposity. Liver is a highly metabolic tissue, which may accumulate fat causing hepatic steatosis. However, when we weighed livers, we found no differences in the Pdk1 KO mice (Fig. 5I). Together, these studies, under the experimental conditions they were conducted in, were unable to confirm that loss of Pdk1 mediates the effect of the *Moo1* locus on obesity (Fig. 5J-M).

**Figure 5.**
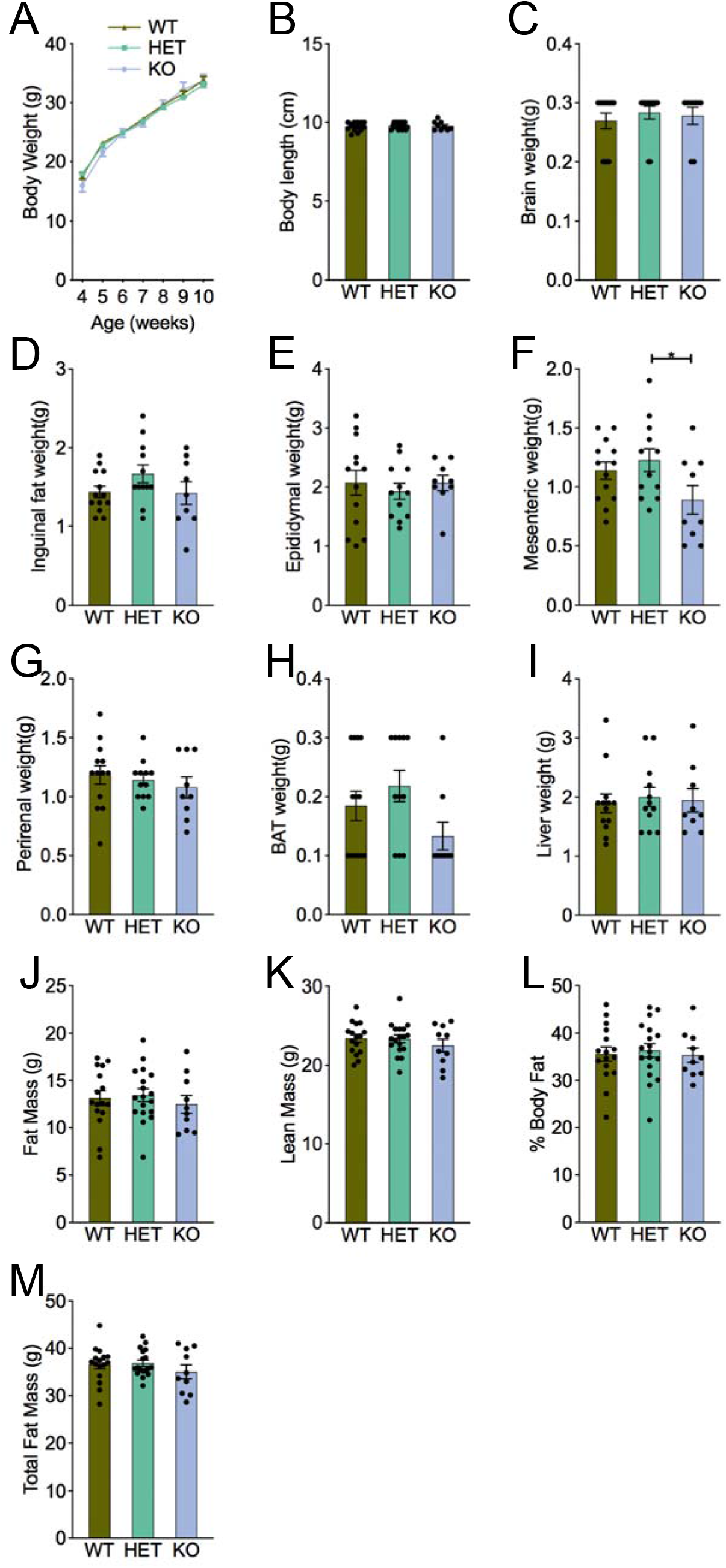
Effect of *Pdk1* on body weight, composition and tissue weight. A.) *Pdk1* Knockout (KO), Heterozygous (HET), and Wildtype (WT) littermate mice were assessed weekly for body weight from 4-10 weeks of age. n=6-14. At 16 weeks *Pdk1* mice were sacrificed and were measured for their (B) body length, (C) brain weight, (D) inguinal fat, (E) epididymal fat, (F) mesenteric fat, (G) perirenal fat, (H) brown adipose tissue (BAT), and (I) liver weight. KO n=9, HET n=18, WT n=16. Body composition analysis in 16 weeks old mice measuring (J) fat mass, (K) lean mass, (L) % body fat and (M) total mass, did not differ between KO, HET, and WT mice. KO n=10, HET n=18, WT n=16. Data were expressed as mean ± SEM and analyzed using one-way ANOVA (p-value shown). Tukey’s pairwise comparisons between groups that were significantly different are shown (*p<0.05).

### Effects of reduced Pdk1 on lipid metabolism

Studies have shown that reducing other isoforms of PDKs may contribute to changes in lipid metabolism and may differ in fed and fasted states (Jeoung et al. 2012; Wu et al. 2018; Jeoung and Harris 2008). However, the *in vivo* effects of PDK1 loss lipid metabolism were not known. We performed analysis of lipid clearance and triglyceride synthesis in *Pdk1* KO mice to assess lipid metabolism. We analyzed the effects of reduced Pdk1 on metabolites in both fasted and non-fasted states. Plasma and liver triglyceride and cholesterol levels did not differ in mice lacking PDK1 (Fig 6A-J). While heart triglyceride levels were unaffected (Fig 6L), heart cholesterol levels were significantly increased, and we observed cardiac lipid accumulation in mice lacking PDK1 (Fig 6K). Collectively, these observations reveal a potential novel role for PDK1 in cardiac lipid metabolism.

**Figure 6.**
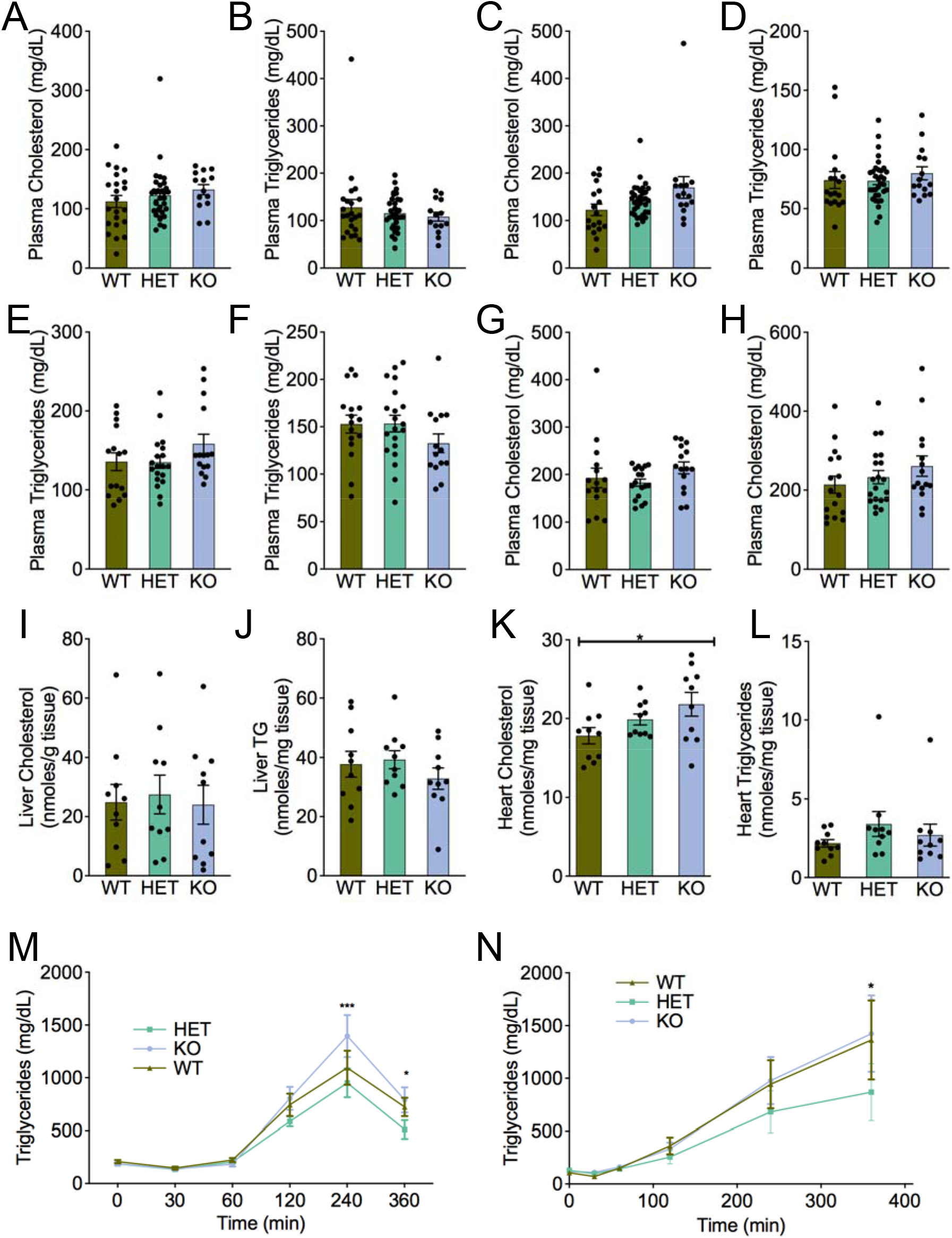
Lipid levels in fasted and non-fasted *Pdk1* mice. Plasma cholesterol levels were measured in (A) non-fasted mice at 6 weeks, (C) fasted mice at 8 weeks, (G) non-fasted mice at 16 weeks, and (H) fasted mice at 16 weeks. Plasma triglyceride levels were measured in (B) non-fasted mice at 6 weeks, (D) fasted mice at 8 weeks, (E) non-fasted mice at 16 weeks, and (F) fasted mice at 16 weeks. n=14-30. Liver (I) cholesterol and (J) triglyceride and heart (K) cholesterol and (L) triglyceride were measured in *Pdk1* KO, HET and WT mice. n=10 in each group. M.) Oral lipid tolerance tests were performed on 14 week old *Pdk1* KO, HET and WT mice. n=12-20. N.) Triglyceride appearance in the plasma during fasting, following lipase inhibition with poloxamer 407 was measured in 12 weeks old mice. n=12-18. Data are expressed as mean ± SEM and were analyzed using repeated measures ANOVA with genotype and time as factors (p-value shown). Tukey’s pairwise comparisons between groups at each time point that were significantly different are shown (*p<0.05, ***p<0.001, for HET vs KO).

### Effects of reduced Itga6 on body weight

Integrins are a family of cell adhesion molecules comprised of a heterodimer of an α- and β-subunit. Integrin alpha 6 (ITGA6) binds with either the β1 or β4 subunits to form heterodimers, which are involved in cell-to-cell adhesion and cell adhesion to the extracellular matrix (Giancotti and Ruoslahti 1999). ITGA6 is known to affect neuronal development and patterning (Marchetti et al. 2013; Pellegatta et al. 2013). Global *Itga6* knockout mice show disrupted neurulation and axial development(Lallier, Whittaker, and DeSimone 1996). Altered neuronal development in areas controlling food intake and/or affecting energy expenditure could theoretically lead to obesity. Thus, *Itga6* is also both a functional and positional candidate gene for mediate the obesity effects at the *Moo1* locus. Based on the a priori knowledge, we hypothesized that reduced ITGA6 activity would promote obesity. We tested this hypothesis using mice with partial deletion of *Itga6* mice. Moo1V^BTBR/BTBR^ mice were heavier than the Moo1V^B6/B6^ mice and the Moo1V^BTBR/BTBR^ had a 50% reduced *Itga6* gene expression. Thus, we hypothesized the *Itga6* HET mice would have an increase in body weight.

We used heterozygous knockout mice in our studies because a homozygous KO *Itga6* mouse is not viable, dying shortly after birth^261^. To determine if reduced Itga6 affects body weight, we measured growth curves as in our previous studies. HET KO mice growth curves were not significantly different compared to their WT, FLOX or CRE littermates. To ensure differences in adiposity were not missed, e.g., due to differences in body size (length) or changes in only specific fat pads, we also directly assessed body fat. In contrast to our expectations, we did not detect differences in body fat measured by DEXA, nor differences in the individual fat pad weights (Fig. 7). These data potentially identify that a global 50% reduction of *Itga6* alone is insufficient to modify body weight, at least using this Cre model under the conditions we studied. An important caveat of these studies is that despite using a strong, ubiquitous expressing cre, at least in the liver in our studies, Itga6 expression was we were unable to detect a significant reduction of *Itga6* in our HET KO mice compared to their controls.

**Figure 7.**
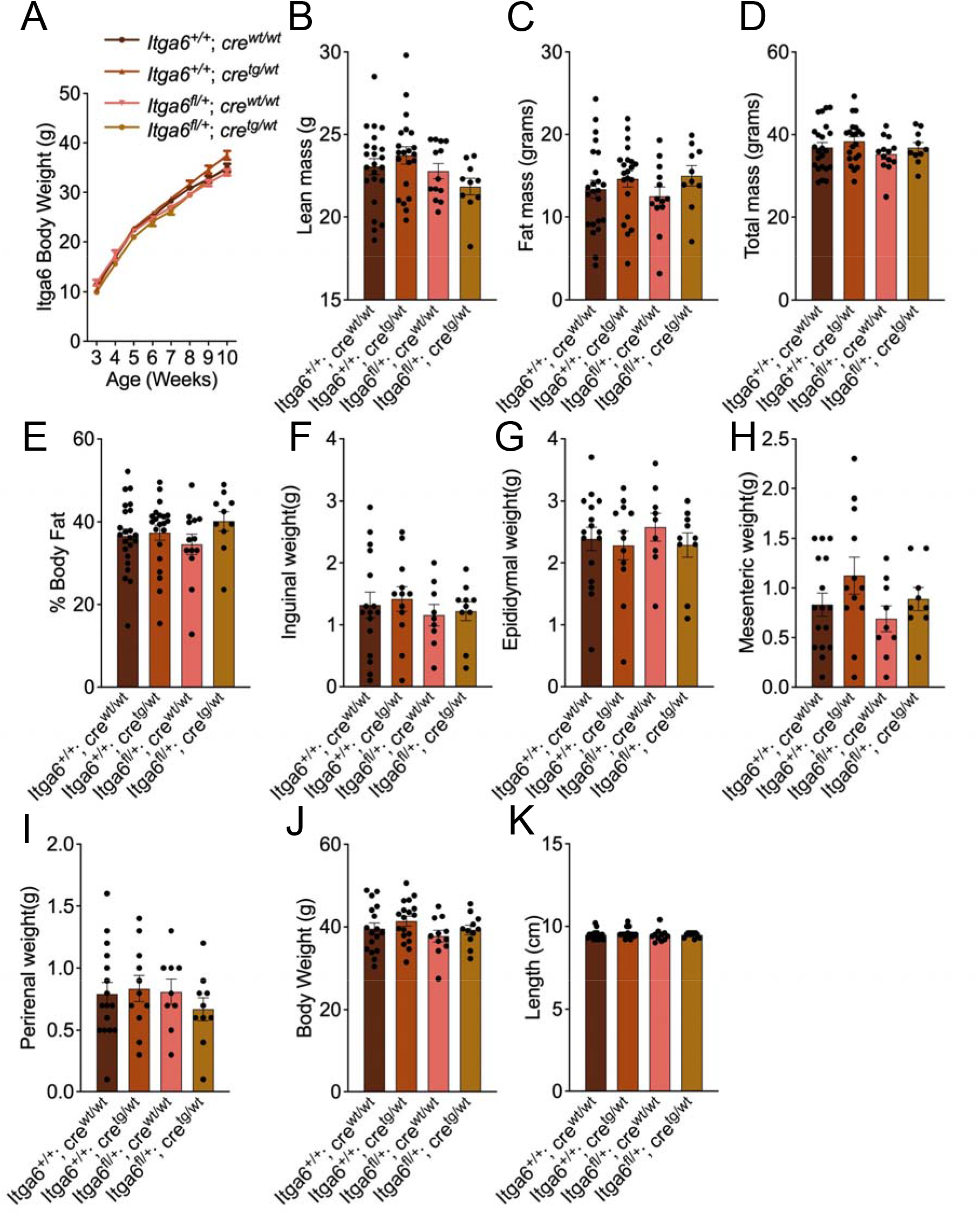
Effect of *Itga6* on body weight, composition, and tissue weight. A.) Growth curves of heterozygous knockout *Itga6*^fl/wt^;Cre^tg/wt^ (HET KO), *Itga6*^wt/wt^;Cre^tg/wt^ (CRE), flox positive *Itga6*^fl/fl^;Cre^wt/wt^ (FLOX), and wildtype *Itga6*^wt/wt^;Cre^wt/wt^ (WT) mice, graphed as mean weekly body weight until 10 weeks of age, as assessed in our congenic studies. n=21-35. (B) Lean mass, (C) fat mass, (D) total fat mass and (E) % body fat (F) Inguinal, (G) Epididymal, (H) Mesenteric and (I) Perirenal fat pad weights, (J) body weight, and (K) body length were measured in 12 weeks old *Itga6* KO and control animals. Data are expressed as mean ± SEM and were analyzed using one-way ANOVA (p-value shown).

## Discussion

The goal of the present study was to elucidate the causal genes that contribute to obesity at the Moo1 QTL. Unfortunately, despite conducting *in vivo* genetic loss-of-function studies on the two leading candidate genes, *Pdk1* and *Itga6*, we were unable to unambiguously point to a single causal gene that mediates the obesity phenotype. Nevertheless, a QTL affecting body weight was localized to the region contained in the Moo1V strain (*Moo1a*). Our inability to identify the mechanism by which this locus affects obesity could have many explanations. The effects of this locus on obesity appear to be susceptible to modulation by the environment (i.e. stress and diet), which likely complicate our efforts to identify the causal gene. It is also possible that both genes must be reduced to see effects on obesity. Another possibility is that a SNP in the *Moo1* region may have long-range interaction that affect a gene or genes outside the locus. There is precedent for variants affecting genes even 1 Mb away (Rask-Andersen, Almén, and Schiöth 2015; Smemo et al. 2014). One of the ESTs, perhaps a directly adjacent gene, *Rapgef4*, or an unknown element causes phenotype; however, there are no known miRNAs in region. More studies will be needed to elucidate this mechanism.

Moo1 QTL has multiple loci: one in region of Moo1V (*Moo1a*), at least one in region of the Moo1U strain, that is most likely located within the Moo1G strain but not within the Moo1L strain (*Moo1b*). This region extends from rs13476570 (top of grey zone for Moo1G) to rs27982530 (top of known B6 region in the Moo1L strain). This ~1.8 Mb region where *Moo1b* is likely located is largely identical by descent between B6 and BTBR mice, with only 111 variants (of 55349 known SNPs and indels) listed as polymorphic between these two strains (retrieved from https://phenome.jax.org/snp/retrieval), none of which are annotated to affect protein coding regions or splice sites. This is markedly in contrast to the 3416 (of 9593 known) variants retrieved from the 316 kb region containing *Moo1a* that differ between B6 and BTBR. Interestingly, *Moo1b* contains part of *Rapgef4*, also known as Epac2, which regulates cAMP signalling and is another potential candidate gene. Given the larger effect size, we focussed on *Moo1a*.The Moo1T strain, which contains *Rapgef4*, does not have a clear obesity phenotype. It is possible that SNPs in Moo1T and not in Moo1V affect this gene causing effect on body weight in opposite direction.

In the current studies, we found that housing environment modified the obesity phenotype of *Moo1*. We found potential suggestive evidence that stress may differentially affect the alleles of *Moo1a*. An increase in stress would counteract the increased adiposity associated with the inheritance of BTBR alleles in this region. Preliminary findings also suggested that the orexin pathway may be involved in the interaction between stress and chromosome 2 genotype. Interestingly, *Pdk1* and *Itga6* have been implicated in the orexin pathway and response to restraint stress, respectively (Gao et al. 2007; Liu, Zhao, and Guo 2016; Orsetti et al. 2008; Sikder and Kodadek 2007; Wan et al. 2017), providing additional support for the findings of these studies.

The significant positive finding from this work was that reduced Pdk1 affects cardiac lipid metabolism. PDK1 inhibitors have been proposed as potential cancer drugs (Sutendra and Michelakis 2013; Zhang et al. 2018) and consistent with our findings, the PDK inhibitor AZ10219759 was reported to cause lipid accumulation in the heart with subsequent necrosis and atrial tissue degeneration (Jones et al. 2014). *Pdk1* may have metabolic effects in the heart that should be considered when using Pdk1 inhibitors as a potential therapeutic. Further studies are needed to understand the role of PDK in cardiac lipid accumulation, ideally using cardiomyocyte-specific knockout mice.

The studies conducted in this paper highlight the importance, and the challenge of physiological studies in mice. It is important to make note of when there are noise windows, construction or facility changes during experiments. There are changes within the facility and environmental factors are hard to control in mouse studies, but may have significant and profound effects on phenotypes and the ability to reproduce phenotypes. Despite these challenges, these studies showed that PDK1 may be important for lipid metabolism and heart function.

## Acknowledgements

We thank Alan Attie for his comments on this manuscript. We thank Elisabeth Georges-Labouesse, Michel Labouesse, and Adele D’Arcangelis for sharing the *Itga6* mice. We also would like to thank the many undergraduate researchers from the Clee lab who contributed to this work.

---

**Supplementary Table 1: Primer sequences**

**Supplementary Figure Legends**

**Supplementary Fig. 1. Genetic Mapping in new F2 B6 × BTBR population.** Body weight (**A**) at 8 weeks of age (blue) and 10 weeks of age (black), and expression of *Itga6* (**B**) and *Pdk1* (**C**) in adipose tissue (green), kidney (magenta), hypothalamus (red), liver (blue), islet (black) and gastrocnemius muscle (purple). Genome positions in Mb on chromosome 2 are shown on the bottom x-axis, and corresponding cM positions are shown along the top. Dashed line indicates genome-wide significance. Small red dot indicates the position of the Moo1-V congenic insert. The peak linkage positions of *Itga6* and *Pdk1* were at 71.694 Mb and 71.74 Mb, respectively. Data were obtained from diabetes.wisc.edu, and are from the study described by Stoehr, J. P et. al (Stoehr et al. 2004b).

**Supplementary Fig. 2. Alignment of ITGA6 protein sequences.** Alignment of ITGA6 protein sequences from vertebrates including mammals, reptiles, fish, birds and amphibians. The amino acid changed to V in the BTBR strain is highlighted and boxed in red. The accession numbers and species identity of the sequences used are as follows: NP_001264899.1 (Mus musculus), XP_002934665.3 (Xenopus tropicalis), NP_001073286.1 (Homo sapiens), XP_006234413.1 (Rattus norvegicus), XP_015145127.2 (Gallus gallus), XP_005202398.1 (Bos Taurus), XP_020931738.1 (Sus scrofa), XP_014869274.1 (Poecilia Mexicana), XP_023416789.1 (Cavia porcellus), XP_006636569.2 (Lepisosteus oculatus), XP_005393272.1 (Chinchilla lanigera), XP_010375889.1 (Rhinopithecus roxellana), XP_011608007.1 (Takifugu rubripes), XP_017591104.1 (Corvus cornix cornix), XP_010601127.1 (Loxodonta Africana), XP_008685728.1 (Ursus maritimus), XP_014455063.2 (Alligator mississippiensis), XP_007431144.1 (Python bivittatus), XP_004032841.1 (Gorilla gorilla gorilla), XP_003640224.1 (Canis lupus familiaris). Alignment was performed using Clustal Omega v 1.2.4.

**Supplementary Fig. 3. Alignment of PDK1 protein sequences.** Alignment of PDK1 protein sequences, including mammals, reptiles, fish, birds and amphibians. The 2 amino acids deleted in the BTBR strain are highlighted and boxed in red. The accession numbers and species names of the sequences used are as follows: NP_766253.2 (Mus musculus), NP_001085016.1 (Xenopus laevis), NP_002601.1(Homo sapiens), NP_446278.2 (Rattus norvegicus), NP_001026523.2 (Gallus gallus), NP_001192886.1(Bos Taurus), NP_001153080.1 (Sus scrofa), XP_014868810.1 (Poecilia Mexicana), XP_003478653.1 (Cavia porcellus), XP_006636568.1 (Lepisosteus oculatus), XP_005393266.1 (Chinchilla lanigera), XP_010375891.1 (Rhinopithecus roxellana), XP_003961887.1 (Takifugu rubripes), XP_010405070.1 (Corvus cornix cornix), XP_010077032.1 (Pterocles gutturalis), XP_010601125.1 (Loxodonta Africana), XP_008685344.1 (Ursus maritimus), XP_006020858.1 (Alligator sinensis), XP_007431139.1 (Python bivittatus), XP_004032847.1 (Gorilla gorilla gorilla), XP_534032.3 (Canis lupus familiaris). Alignment was performed using Clustal Omega v 1.2.4.

**Supplementary Fig. 4. Strain distribution pattern of SNPs affecting amino acids in ITGA6 and PDK1.** Data were obtained from the Mouse Phenome Database, Sanger4 dataset (www.jax.org/phenome). In both cases, BTBR appears to have the ancestral allele.

**Supplementary Fig. 5. Environmental modulation of body weight phenotypes in the Moo1V strain.** [V strain growth curves CDM rm 1 vs 2 vs 3; note that panel 1 is same data as in fig X; also show AT weights in CDM rm 3] (A) Body weights of Moo1V strain mice housed in CDM room1 weighed weekly from 3 to 10 weeks of age n=11-21) (B) Body weights of Moo1V strain mice housed in CDM room 2 weighed weekly from 3 to 10 weeks of age. n=33-45. (C) Body weights of Moo1V strain mice housed in CDM room 3 weighed weekly from 3 to 10 weeks of age. n=20-39. Genotype shown on graph refers to genotype in congenic region of the Moo1V strain. (D) % body fat was measured by DEXA in 12 weeks old Moo1V strain mice. n= 29 for BTBR/BTBR and B6/B6 and n=30 for B6/BTBR Moo1V genotypes. Data are shown mean ± SEM and were analyzed by repeated measures ANOVA with weeks, age and genotype as the factors. Tukey’s pairwise comparisons between groups that were significantly different were shown (*p<0.05).

