## Supplemental Figure 2 for "Analysis of a genetic region affecting mouse body weight"

| fugu  shortfin_molly  gar  frog  elephant  bear  dog  cow  pig  guinea_pig  chinchilla  human  gorilla  monkey  mouse  rat  python  alligator  chicken  crow | MDSSTMLLHGRNFAGLYLLSYLLLPTSVLTFNLDT—-SHTIRKDGDPGSLFGFSLAMHRQ  -----MEGNDTTRGLRLLALLLACGGLLSAFNLDT—-EDVLRRDGDPGSLFGFSLAMHRQ  -----MAFSATSFFISFSLICLLKRTHVSAFNLDT—-VNVLRRNGEPGSLFGFSLALHRQ  -----MRLAA---ARINLLLG-CCCAFIAAFNLDT--ENVISKSGDPGSLFGFSMAMHWQ  -----MAAAGQL-WLLYLSAG-LLSLLCAAFNLDTREDYVIQKLGDPESLFGFSLAMHWQ  -------------------------------------------------MVADEAVRGES  -----MAAAGRL-WLLYLSAG-LLPRLGAAFNLDTREDHVIRKAGEPGSLFGFSLAMHRQ  -----MAAAGRL-WLLYLSVG-LLPRLGTAFNLDTREDNVIEKTGDSGSLFGFSLAMHWQ  -----MAAAGRL-WLLYLSAG-LLPRLGAAFNLDTREDNVIEKTGDSGSLFGFSLAMHRQ  ------------------------------------------------------------  -----MAAAGQL-CLLYLSAV-LLPRLGAAFNLDTREDNVIRKSGDPGSLFGFSLAMHWQ  -----MAAAGQL-CLLYLSAG-LLSRLGAAFNLDTREDNVIRKYGDPGSLFGFSLAMHWQ  -----MAAAGQL-CLLYLSAG-LLSRLGAAFNLDTREDNVIRKYGDPGSLFGFSLAMHWQ  -----MAAAGQL-CLLYLSAG-LLPRLGSAFNLDAEEDNVIRKYGDPGSLFGFSLAMHWQ  -----MAVAGQL-CLLYLSAG-LLARLGTAFNLDTREDNVIRKSGDPGSLFGFSLAMHWQ  -----MAVAGQL-CLLYLSAG-LLARLSTAFNLDTREDNVIRKSGDPGSLFGFSLAMHWQ  -------------MGSLFYLC-LLPGIVRAFNLDT--ENVVKWNGEPGSFFGFSLAMHQQ  -----MAAPLGLYLRLGQWLL-LLPGLARAFNLDA--EQAVVRRGGKGSLFGFSLAMHWQ  -----MAA------ALLLYLP-LLPGLAGAFNLDA--ENVIGRRGEPGSLFGFSLAMHRQ  ------------------------------------------------------------ | 58  53  53  49  53  11  53  53  53  0  53  53  53  53  53  53  44  52  46  0 |
| --- | --- | --- |

| fugu  shortfin_molly  gar  frog  elephant  bear  dog  cow  pig  guinea_pig  chinchilla  human  gorilla  monkey  mouse  rat  python  alligator  chicken  crow | INPD-KRILLIGAPRAKALGKQTSSFTGGLYKCEITP-FKDCERVDFDNEENTQTEKKEN  LHPEDKRILLVGAPQAKALNGQKSNVTGGIYKCEMSSNSDACSRVVFDNNEDSENENKEN  LHPAEKRILLVGSPRAKALKNQKANVTGGLYSCEISPQLGDCKRIEFDNEADLSSESKEN  LDPVNRRLLLVGSPRAKALPAQKANVTGGLYVCDITSPSPSCNRVLFDDSVDLTSESKEN  LQPEDKRLLLVGAPQAKALPLQRANRTGGLYSCDITS-QEPCTRIEFDNDADPSSESKED  AQKSYSERLLVGAPRAAALPLQRANRTGGLYSCDITS-PGPCTRIEFDNDADPSSESKED  LQPEDRRLLLVGAPRAVALPLQRANRTGGLYSCDITS-PGPCTRIEFDNDADPSSESKED  LQPEDKRLLLVGAPRAVALPLQKANRTGGLYSCDITS-RGPCTRIEFDNDADPSSESKED  LEPEEKRLVLVGAPRAVALPLQKANRTGGLYSCDITS-RGPCTRIEFDNDADPSSESKED  ------------------------------------------------------------  LHPEDKRLLLVGAPRAKALPRQRANRTGGLYSCDITS-RGPCSRVEFDNDADPASESKED  LQPEDKRLLLVGAPRAEALPLQRANRTGGLYSCDITA-RGPCTRIEFDNDADPTSESKED  LQPEDKRLLLVGAPRAEALPLQRANRTGGLYSCDITA-RGPCTRIEFDNDADPTSESKED  LQPEDKRLLLVGAPRAEALPLQRANRTGGLYSCDITA-RGPCTRIEFDNDADPTSESKED  LQPEDKRLLLVGAPRAEALPLQRANRTGGLYSCDITS-RGPCTRIEFDNDADPMSESKED  LQPEDKRLLLVGAPRAEALPLQKANRTGGLYSCDITS-RGPCTRIVFDNDADPMSESKED  LDPN-RRLLLVGAPRQKAFPMQRANRTGGLFSCDIETPDLPCTRIEFDETADLAKESKED  LQPQDKRLLLVGAPREKAFPAQQANRTGGLYWCDIIS-ETYCTRVMFDEKTDPEKESKED  LQPQEKRLLLVGAPREKAFPSQQANRTGGLYSCDITSSDTRCTRVVFDEDTDPKMESKED  ------------------------------------------MRVKFDEETDPKMESKED | 116  113  113  109  112  70  112  112  112  0  112  112  112  112  112  112  103  111  106  18 |
| --- | --- | --- |

| fugu  shortfin_molly  gar  frog  elephant  bear  dog  cow  pig  guinea_pig  chinchilla  human  gorilla  monkey  mouse  rat  python  alligator  chicken  crow | QWMGVMVQSQGPGGKIVTCAHRYQRRMFANTPQESWDITGRCYVLSQDLTIDVSSDEDGG  QWMGVTVSSQGPGGKVLTCAHRYKRQRNVNSPHESRDIIGRCYVLSQDLTIDPSSSEDGG  QWMGVTVQSQGPGGKIVTCAHRYQRRMFVGSAQESRDITGRCYVLSENLQINEGSEEDGG  QWMGVSVQSQGPGGKIVTCAHRYERRTSVNTNYENRDITGRCYVFSQDLTMTD--EMDGG  QWMGVTVQSQGPGGKVVTCAHRYEKRQHVNTKQESRDIFGRCYVLSQNLRIED--DMDGG  QWMGVTVQSQGPGGKIVTCAHRYEKRQHVNTKQESRDIFGRCYVLSQNLRIED--DMDGG  QWMGVTVQSQGPGGKVVTCAHRYEKRQHVNTKQESRDIFGRCYVLSQNLRIED--DMDGG  QWMGVTVQSQGPGGKVVTCAHRYEKRQHVNTKQESRDIFGRCYVLSQNLRIED--DMDGG  QWMGVTVQSQGPGGKVVTCAHRYEKRQHVNTKQESRDIFGRCYVLSQNLRIED--YMDGG  --MGVTVQSQGPGGKVVTCAHRYEKRQHVNTKQESRDIFGRCYVLSQNLRIED--DMDGG  QWMGVTVQSQGPGGKVVTCAHRYEKRQHVNTKQESRDIFGRCYVLSQNLKIED--DMDGG  QWMGVTVQSQGPGGKVVTCAHRYEKRQHVNTKQESRDIFGRCYVLSQNLRIED--DMDGG  QWMGVTVQSQGPGGKVVTCAHRYEKRQHVNTKQESRDIFGRCYVLSQNLRIED--DMDGG  QWMGVTVQSQGPGGKVVTCAHRYEKRQHVNTKQESRDIFGRCYVLSQNLRIED--DMDGG  QWMGVTVQSQGPGGKVVTCAHRYEKRQHVNTKQESRDIFGRCYVLSQNLRIED--DMDGG  QWMGVTVQSQGPGGKVVTCAHRYEKRQHVNTKQESRDIFGRCYVLSQNLRIED--DMDGG  QWMGVTVRSQGPGGKVVTCAHRYEKRYYVNTEQETRDIIGRCYVLSQDLSISNNEYMDGG  QWMGVTVRSQGPGGKVVTCAHRYEKRQYVNTLQETRDIIGRCYVLSQNLTIVD--DMDGG  QWMGVTVQSQGPGGNVVTCAHRYEKRQYVNTVQETRDIIGRCYVLSQDLTIKD--DMDNG  QWMGVTVQSQGPGGNVVTCAHRYEKRQYVNTVQETRDIIGRCYVLSQDLTIKD--DMDNG | 176  173  173  167  170  128  170  170  170  56  170  170  170  170  170  170  163  169  164  76 |
| --- | --- | --- |
| fugu  shortfin_molly  gar  frog  elephant  bear  dog  cow  pig  guinea_pig  chinchilla  human  gorilla  monkey  mouse  rat  python  alligator  chicken  crow | NWNFCDGRPRGHEMFGSCQQGLAATFTKDYHYLVFGAPGAYNWKGIVRMEQKNNTLLEMG  SWHFCNNRNRGHERFGSCQQGLSATFDKDFHYFIFGAPGAYNWKGVVRLEQKNDTLIDMG  NWKFCEGRTRGHERFGSCQQGLSATFTKDYHYVVFGAPGAYNWKGIVRVEQKNSTLVDLG  DWNFCEGRRRGHEQFGSCQQGVAATFTKDFHYIVFGAPGTYNWKGVVRAEQKNNTLEDLF  DWSFCDGRLRGHEKFGSCQQGVAATFTKDFHYIVFGAPGTYNWKGIVRVEQKNNTFFDMN  DWSFCDGRLRGHEKFGSCQQGVAATFTKDFHYIVFGAPGTYNWKGIVRVEQKNNTFFDMN  DWSFCDGRLRGHEKFGSCQQGVAATFTKDFHYIVFGAPGTYNWKGIVRVEQKNNTFFDMN  DWSFCDGRLRGHEKFGSCQQGVAATFTKDFHYIAFGAPGTYNWKGIVRVEQKNNTFFDMN  DWSFCDGRLRGHEKFGSCQQGVAATFTKDYHYIVFGAPGTYNWKGIVRVEQKNNTFFDMN  DWSFCDGRLRGHEKFGSCQQGVAATFTKDFHYIVFGAPGTYNWKGIVRVEQKNNTFFDMN  DWSFCDGRLRGHEKFGSCQQGVAATFTKDFHYIVFGAPGTYNWKGIVRVEQKNNTFFDMN  DWSFCDGRLRGHEKFGSCQQGVAATFTKDFHYIVFGAPGTYNWKGIVRVEQKNNTFFDMN  DWSFCDGRLRGHEKFGSCQQGVAATFTKDFHYIVFGAPGTYNWKGIVRVEQKNNTFFDMN  DWSFCDGRLRGHEKFGSCQQGVAATFTKDFHYIVFGAPGTYNWKGIVRVEQKNNTFFDMN  DWSFCDGRLRGHEKFGSCQQGVAATFTKDFHYIVFGAPGTYNWKGIVRVEQKNNTFFDMN  DWSFCDGRLRGHEKFGSCQQGVAATFTKDFHYIVFGAPGTYNWKGIVRVEQKNNTFFDMN  EWSFCDGRLRGHEKFGSCQQGVAATFTRDNHYIVFGAPGTYNWKGVVRAEQKNHTLDEMG  EWSFCDGRLRGHEKFGSCQQGVAATFTRDYHYIVFGAPGTYNWKGVVRAEQKNQTFYELS  VWSFCDGRLRGHEKFGSCQQGVAATFTRDYHYIVFGAPGTYNWKGVVRAEQKNQTFYDLG  VWSFCDGRLRGHEKFGSCQQGVAATFTRDYHYIVFGAPGTYNWKGVVRAEQKNQTFYDLG | 236  233  233  227  230  188  230  230  230  116  230  230  230  230  230  230  223  229  224  136 |

| fugu  shortfin_molly  gar  frog  elephant  bear  dog  cow  pig  guinea_pig  chinchilla  human  gorilla  monkey  mouse  rat  python  alligator  chicken  crow | VYDDGPYEVGDEHLLNPDLVPLPANSYLGFSLDTGHRVISSLVLTVVAGAPRANHSGAVV  IYDDGPFEAGDETEKKPDLVPAPPNSYMGFSLDSGKALTNKGQLTVVAGAPRAYFSGGVI  IYEDGPYEVGDETQFDANLVPVPDNSYLGFSLDSGHNLIQKGQLTIVSGAPRAKHSGAVV  IFEDGPYETGGENKRDSDLVPAPDNSYLGFSLDTGRGITSKEEMTFVSGAPRANHSGAVV  IFEDGPYEVGGENNHDESLVPVPANSYLGFSLDSGKGIVSKDDITFVSGAPRANHSGAVV  IFEDGPYEVGGETDHDESLVPVPANSYLGFSLDSGKGIVSKDEITFVSGAPRANHSGAVV  IFEDGPYEVGGETDHDESLVPVPANSYLGFSLDSGKGIVSKDEITFVSGAPRANHSGAVV  IFEDGPYEVGGETDHDESLVPVPANSYLGFSLDSGKGIVSKDEITFVSGAPRANHSGAVV  IFEDGPYEVGGETDNDESLVPVPANSYLGFSLDSGKGIVSKDEITFVSGAPRANHSGAVV  IFEDGPYEVGGETDHDESLVPVPANSYLGFSLDSGKGIVSKDEITFVSGAPRANHSGAVV  IFEDGPYEVGGETDHDESLVPVPANSYLGFSLDSGKGIVSKDEITFVSGAPRANHSGAVV  IFEDGPYEVGGETEHDESLVPVPANSYLGFSLDSGKGIVSKDEITFVSGAPRANHSGAVV  IFEDGPYEVGGETEHDESLVPVPANSYLGFSLDSGKGIVSKDEITFVSGAPRANHSGAVV  IYEDGPYEVGGETEHDESLVPVPANSYLGFSLDSGKGIVSKDEITFVSGAPRANHSGAVV  IFEDGPYEVGGETDHDESLVPVPANSYLGFSLDSGKGIVSKDDITFVSGAPRANHSGAVV  IFEDGPYEVGGETDHDESLVPVPANSYLGFSLDSGKGIVSKDDITFVSGAPRANHSGAVV  IFDDGPYEVGGESLRKETLVPVPANSYLGFSLDSGKGIVSQEDITFVSGAPRANHSGAVV  IFDDGPYEVGDEASRDESLVPVPANSYLGFSLDSGKGIVSQEEITFVSGAPRANHSGAVV  IFDDGPYEVGDESRQDKNLVPVPANSYLGFSLDSGKGIVSQDEMTFVSGAPRANHSGAVV  IFDDGPYEVGDESHQDKNLVPVPANSYLGFSLDSGTGIVSQDEMTFVSGAPRANHSGAVV | 296  293  293  287  290  248  290  290  290  176  290  290  290  290  290  290  283  289  284  196 |
| --- | --- | --- |

| fugu  shortfin_molly  gar  frog  elephant  bear  dog  cow  pig  guinea_pig  chinchilla  human  gorilla  monkey  mouse  rat  python  alligator  chicken  crow | LLKKKSDASFKLVVEHTFYGPGLASSFGYDVAVVDLNGDGWQDIVVGAPQFYMK-DGDVG  LLKKGGELSRILEKEYILKGEGLASSFGYDLTVLDLNGDGWDDIVVGAPQYFEK-DSEIG  LLKTGGTESRVLNKEFVLEGPGLASSFGYDVAVVDLNSDGWQDIVVGAPQYFVK-DGETG  LLQKFPPNERMLSAVHTFQGEGLASSFGYDVAVVDLNRDGWKDIVVGAPQYFDKNDREIG  LLKRDMK-STHLLPEHIFDGEGLASSFGYDVAVVDLNKDGWQDIVIGAPQYFDR-GGEVG  LLKRDLK-SAHLLPEYIFDGEGLASSFGYDVAVADLNKDGWQDIIVGAPQYFDR-DGEVG  LLKRDMK-SAHLLPEYIFDGEGLASSFGYDVAVADLNKDGWQDIVIGAPQYFDR-DGEVG  LLKRDLK-SAHLLPEHIFDGEGLASSFGYDVAVVDLNKDGWQDIVIGAPQYFDR-SGEVG  LLKRDLK-SAHLLPEHIFDGEGLASSFGYDVAVVDLNSDGWQDIVIGAPQYFDR-SGEVG  LLKRELK-SAHLLPEHIFDGEGLASSFGYDVAVVDLNKDGWQDIVIGAPQYFDR-DGEIG  LLKRELK-SAHLLPEHIFDGEGLASSFGYDVAVVDLNKDGWQDIVIGAPQYFDR-DGEIG  LLKRDMK-SAHLLPEHIFDGEGLASSFGYDVAVVDLNKDGWQDIVIGAPQYFDR-DGEVG  LLKRDMK-SAHLLPEHIFDGEGLASSFGYDVAVVDLNKDGWQDIVIGAPQYFDR-DGEVG  LLRRDMK-SAHLLPEHIFDGEGLASSFGYDVAVVDLNKDGWQDIVIGAPQYFDR-DGEVG  LLKRDMK-SAHLLPEYIFDGEGLASSFGYDVAVVDLNADGWQDIVIGAPQYFDR-DGEVG  LLKRDMK-SAHLLLEYIFDGEGLASSFGYDVAVVDLNADGWQDIVIGAPQYFDR-DGEVG  LLKKDSDVQRALSPVHMFEGEGLASSFGYDVAVVDLNSDGWQDIVVGAPQYFDR-DGEIG  LLRKDPSIQSALSLVHIFEGEGLASSFGYDVAVVDLNKDGWQDIVVGAPQYFER-DEDIG  LLKKEK-NQRALSLEHMFEGEGLASSFGYDVAVVDLNSDGWQDIVVGAPQYFDR-SGDIG  LLKK-K-NQRALSLEHMFEGEGLASSFGYDVAVVDLNSDGWQDIVVGAPQYFDR-SGDIG | 355  352  352  347  348  306  348  348  348  234  348  348  348  348  348  348  342  348  342  253 |
| --- | --- | --- |

| fugu  shortfin_molly  gar  frog  elephant  bear  dog  cow  pig  guinea_pig  chinchilla  human  gorilla  monkey  mouse  rat  python  alligator  chicken  crow | GAAFVYMNREGRWDGVMPVRLNGTKDSMFGLAVENIGDVNQDSFEDIAVGAPNEEGGAGR  GAVYVYINKAGNWNEVTPTRIDGPAVSMFGLAVENLGDINQDGYHDFAVGAPHEDSGAGK  GAVYVYINKNGKWDQITPIQLNGTKDSMFGLAVENIGDINLDGFQDIAVGAPYDDAGSGK  GAIYVYINQQGNWNDVMPVRIVGTKDSMFGISVKNIGDINQDGYPDIAVGAPYDD-GFGK  GAVYVYINQQGRWNNVKPIRLNGTKDSMFGIAVKNIGDINQDGYPDIAVGAPYDG--MGK  GAVYVYINEHGRWNNVKPIRLNGTKDSMFGVAVKNIGDINQDGYPDIAVGAPYDD--MGK  GAVYVYINQQGRWNNVKPIRLNGTKDSMFGIAVKNIGDINQDGYPDIAVGAPYDD--MGK  GAAYVYINQQGRWNNVKPIRLNGTKDSMFGIAVKNIGDINQDGYPDIAVGAPYDG--KGK  GAVYVYMNQQGRWKNVKPIRLNGTKDSMFGVAVKNIGDINQDGYPDIAVGAPYDG--KGK  GAAYVYINQQGRWNNVKPIRLNGTKDSMFGIAVKNIGDINQDGYPDIAVGAPYDD--MGK  GAVYVYINQQGRWNNVKPIRLNGTKDSMFGIAVKNIGDINQDGYPDIAVGAPYDD--MGK  GAVYVYMNQQGRWNNVKPIRLNGTKDSMFGIAVKNIGDINQDGYPDIAVGAPYDD--LGK  GAVYVYMNQQGRWNNVKPIRLNGTKDSMFGIAVKNIGDINQDGYPDIAVGAPYDD--LGK  GAVYVYMNQQGRWNNVKPIRLNGTKDSMFGIAVKNIGDINQDGYPDIAVGAPYDD--TGK  GAVYVYINQQGKWSNVKPIRLNGTKDSMFGISVKNIGDINQDGYPDIAVGAPYDD--LGK  GAVYVYINQQGRWSNVKPVRLNGTKDSMFGIAVKNIGDINQDGYPDIAVGAPYDD--LGK  GAAYVYINQEGKWDRVKPIRLNGSTDSMFGIAVENVGDVNKDGFPDIAVGAPYDG-DFGK  GAVYIYINQNGNWEGVKPVRLNGTTDSMFGLAVESIGDVNQDGFPDIAVGAPYDG--SGK  GAVYIYINQRGKWEGIKPIRLNGTADSMFGLAVENVGDINQDGYPDIAVGAPYDG--FGK  GAVYIYMNRQGKWAGVKPLRLNGTAGSMFGLAVENVGDINQDGYPDIAVGAPYDG--FGK | 415  412  412  406  406  364  406  406  406  292  406  406  406  406  406  406  401  406  400  311 |
| --- | --- | --- |

| fugu  shortfin_molly  gar  frog  elephant  bear  dog  cow  pig  guinea_pig  chinchilla  human  gorilla  monkey  mouse  rat  python  alligator  chicken  crow | VFIYHGSKQGV-KTRPAQILSGKGYNMKLFGYSLAGNMDLDGNSYPDVAVGSLSDTAVIF  VYIYHGSARGEITDKASQVLSSKSAGVRQFGYSLAGNMDLDKNSYPDLAVGSLSDSVFVY  VYIYHGSKSGI-TTKPAQVLEGRDHNIKLFGYSLAGNMDLDRNSYPDIAIGSLSDSVFVY  VYVYHGSKDGI-ISEPAQVIEGRSTNTRFFGYSIAGDMDLDQNSYPDIAVGSLSDSVKVF  VFIYHGSANGI-NTKPTQVLKGR---SPYFGYSIAGNMDLDRNSYPDVAVGSLSDSVTVF  VFIYHGSPNGI-NTKPTQILEGK---SPYFGYSIAGNMDLDRNSYPDVAVGSLSDSVAVF  VFIYHGSPNGI-NTKPTQILEGK---SPYFGYSIAGNMDLDRNSYPDVAVGSLSDSVTVF  VFIYHGSANGI-NTKPTQVLEGK---SPFFGYSIAGNMDLDRNSYPDVAVGSLSDSVTIF  VFIYHGSANGI-NTKPTQVLEGK---SPSFGYSIAGNMDLDRNWYPDVAVGSLSDSVTIF  VFIYHGSANGI-NTKPTQVLEGT---TPSFGYSIAGNMDLDRNSYPDVAVLP-SQISSIF  VFIYHGSANGI-NTKPTQILEGT---TPYFGYSIAGNMDLDRNSYPDVAVGSLSDSVTIF  VFIYHGSANGI-NTKPTQVLKGI---SPYFGYSIAGNMDLDRNSYPDVAVGSLSDSVTIF  VFIYHGSANGI-NTKPTQVLKGI---SPYFGYSIAGNMDLDRNSYPDVAVGSLSDSVTIF  VFIYHGSANGI-NTKPTQVLKGK---SPYFGYSIAGNMDLDRNSYPDVAVGSLSDSVTIF  VFIYHGSPTGI-ITKPTQVLEGT---SPYFGYSIAGNMDLDRNSYPDLAVGSLSDSVTIF  VFIYHGSPTGI-ITKPTQVLEGT---TPFFGYSIAGNMDLDRNSYPDVAVGSLSDSVTIF  VYLYHGSRNGI-NTKPAQILDGGTNNVIFFGYSITGNMDLDGNSYPDIAVGSLSDSVSVY  VYIYHGSKNGI-NTKATQILDGEENNIQFFGYSIAGNMDLDRNSYPDIAVGSLSDSVSVY  VYIYHGSKNGI-NTEPAQILDGEKTGTNFFGYSIAGNMDLDKNSYPDIAVGSLSDSVSVF  VYIYHGSENGI-NTKPAQILDGEKTNTNFFGYSVAGNMDLDENSYPDVAVGSLSDSVNVY | 474  472  471  465  462  420  462  462  462  347  462  462  462  462  462  462  460  465  459  370 |
| --- | --- | --- |

| fugu  shortfin_molly  gar  frog  elephant  bear  dog  cow  pig  guinea_pig  chinchilla  human  gorilla  monkey  mouse  rat  python  alligator  chicken  crow | RARPIISIQRDVRVSPQEVDLAVK-TCG--NSICFTVDTCFTYTANTASYNPKLTVGFSV  RARPVVNIQKKITFSPSRINLSQK-NCG--NTFCLKVKTCFTYTANPSSYAPRLTVGYSL  RSKPVVNIVKEITITPKDIDLKKPAPCLDGKGFCLTVKACFEYTANPKDYNPRLTMDFTF  RSRPVITIKKTIKVTPDRIDFNKK-NCDAPSGICLEVEGCFEYTANPKTYNPVLTLSCTF  RARPVINIQKTITVTPDKIDLSKKMSCAAPSGICLKVKACFEYTAKPTGYNPSIVIVGTL  RSRPVITIQKTLTVTPGTIDLRQKTFCGAPSGICLKVKACFEYTAKPAGYNPLITILGTL  RSRPVINIQKTVTITPDTIDLRQKTLCGAPSGICLKVKACFEYTAKPTGYNPSISILGTL  RSRPVINIQQTITVTPNRIDLRQKTLCGAPSGICLKVKACFEYTAKPTGYNPSITIVGTL  RSRPVINIQQTITVTPNRIDLRQKTSCGAPSGICLKVKACFEYTTKPANDDPSITIVGTL  RSRPVINIQKTITVTPSRIDLRQK-TCAAPSGICLKVKACFEYTAKPTGYNPPITIVGTL  RTRPVINIQKIITVTPNRIDLRQK-MCRAPSGICLKVKACFEYTAKPTGYSPAITIVGTL  RSRPVINIQKTITVTPNRIDLRQKTACGAPSGICLQVKSCFEYTANPAGYNPSISIVGTL  RSRPVINIQKTITVTPNRIDLRQKTACGAPSGICLQVKSCFEYTANPAGYNPSISIVGTL  RSRPVINIQKTITVTPNRIDLRQKTACGAPSGICLQVKACFEYTAKPAGYNPSISIVGTL  RSRPVINILKTITVTPNRIDLRQKSMCGSPSGICLKVKACFEYTAKPSGYNPPISILGIL  RSRPVINILKTITVTPSRIDLRQKSMCGSPSGICLKVKACFEYTAKPSGYNPPISILGIL  RSRPVINIKKTIMISPDKIDLNMR-NCDVSSNLCLNIRSCFEYTASESNFNQKIILNYSF  RSRPVISIKKTIKILPDKIDLNRR-TCPEPSGICLNVEACFEYTASPQDFNPKIKLNYTF  RSRPVISITKSITVQPDKLDLKKK-NPEDPSEIWMDVKACFQYTANPRNLNPRIKINYTF  RSRPVISIRRNITVQPDRIDLKKK-NPEDPGEIRMDVKACFKYTANPRDLNPRIKINYMF | 531  529  531  524  522  480  522  522  522  406  521  522  522  522  522  522  519  524  518  429 |
| --- | --- | --- |

| fugu  shortfin_molly  gar  frog  elephant  bear  dog  cow  pig  guinea_pig  chinchilla  human  gorilla  monkey  mouse  rat  python  alligator  chicken  crow | ESDGDRRKQGLPSRLVFLNMSRSDTDYQFNGTLDLRSQKQETCIKILGKLKDNIKDKLRS  EADADRRKTNLLPRAIFTEPSDSDRDYIYKGTITMDTKGREQCFTRQLAIQENIKDKLRA  EADSDRRKINLQPRVSFSERKATDPDNQYSGTLELRGQNQKKCIQAEGKLQD-TTDKLRG  ELENDRRLLQKPLRMNFKDLP---QESKLPKTIELRGQNQRRCVTTTLQLLESIRDKLHP  EAEKERRKSGLSSRVQIRNQG---SEPKYSQQLILSRQKQKACMDETLWLQENIRDKLRP  EAEKERRKSGLSSRVHFRNQG---SEAKYTQELTLNRQKQKACIEETLWLQENIRDKLRP  EAEKERRKSGLSSRVQFRNQG---SEPRYTQELTLNRQKQKACMEETLWLQENIRDKLRP  EAEKERRKSGLSSRVQFRNQG---SEPKYSQELTLNRQKQKACMEETLWLQENIRDKLRP  EAEKERRKSGLSSRVQFRNQG---SEPKYTQELTLHRQKQKACMEETLWLQENIRDKLRP  EAEKQRRKSGLSSRVQFRNQA---SEPRYTQELTLQRQKQRLCMEETLWLQENIRDKLRP  EAEKQRRKSGLSSRVQFQNQA---SEPRYTQELTLTRQKQRQCMEETLWLQENIRDKLRP  EAEKERRKSGLSSRVQFRNQG---SEPKYTQELTLKRQKQKVCMEETLWLQDNIRDKLRP  EAEKERRKSGLSSRVQFRNQG---SEPKYTQELTLKRQKQKVCMEETLWLQDNIRDKLRP  EAEKERRKSGLSSRVQFRNQG---SEPKYTQELTLKRQKQKVCMEETLWLQDNIRDKLRP  EAEKERRKSGLSSRVQFRNQG---SEPKYTQELTLNRQKQRACMEETLWLQENIRDKLRP  EAEKERRKSGLSSRVQFRNQG---SEPKYTQELTLNRQKQRACMEETLWLQENIRDKLRP  EAENERRRLGLPPRVHFSKLR---SVP-FSGSVTLRGQKLEECVTTRLELEEKIKDKLRP  EAEKDRRRLGLPSRVHFPNHP---SDQ-LTGSIKLQGQNTRECVKTSLVLQENIKDKLRP  EAENERRQLGLPSRVRFKDYL---SDQ-FTASTTLIGQNSKRCVTAKLVLQEKIKDKLRP  EVENERRQLGLPSRVRFIDHS---SDQ-FTASTTLRGQNSWECVTTKLVLQEKIKDKLRP | 591  589  590  581  579  537  579  579  579  463  578  579  579  579  579  579  575  580  574  485 |
| --- | --- | --- |

| fugu  shortfin_molly  gar  frog  elephant  bear  dog  cow  pig  guinea_pig  chinchilla  human  gorilla  monkey  mouse  rat  python  alligator  chicken  crow | IPIEVSSEILGT-----RRHKSKNGLPQLMPIQDASQLSKDVVMVNFVKEGCGSDHVCNS  IPIDVSVNIQDA-----QRKRRQTQAPQLSPALDANDAQPTRKKVEFIKEGCGNDNVCQS  IPIAVSVAIKNA-----KRRKRQSALAELVPVLSSELPTR--SEVNFVKEGCGTDNICQS  IAVSVSAEIKSV----RRRKRQSSPLPELMPILNSNEPKNATANVQFLKEGCGEDNICNS  IPITASVEIQES-----GSRRRVNSLPEIPPILNSNEPKTVHADVHFLKEGCGSDNVCNS  IPITASVEIQEP-----NTRRRVNSLPEVLPILNSNEPKTVHRDVHFLKEGCGDDNVCNS  IPITASVEIQEP-----NTRRRVNSLPEILPILNSNEPKTVHRDVHFLKEGCGDDNICNS  IPITASVEIQEP-----STRRRVNSLPEVLPILNSNEPKFVHTDVHFLKEGCGDDNICNS  IPITASVEIQEP-----STRRRVNSLPEVLPILNSNEPKSVHTDVHFLKEGCGADNICNS  IPITASVEIQEP-----SSRRRVNSLPEILPILNSNEPTTVQTDVHFLKEGCGDDNVCNS  IPITASVEIQEP-----SSRRRVNSLPEILPILNSNEPKTIQTDVHFLKEGCGDDNVCNS  IPITASVEIQEP-----SSRRRVNSLPEVLPILNSDEPKTAHIDVHFLKEGCGDDNVCNS  IPITASVEIQEP-----SSRRRVNSLPEVLPILNSDEPKTAHIDVHFLKEGCGDDNVCNS  IPITASVEIQEP-----SSRRRVNSLPEVLPILNSDEPKTAHTDVHFLKEGCGDDNVCNS  IPITASVEIQEP-----SSRRRVNSLPEVLPILNSNEAKTVQTDVHFLKEGCGDDNVCNS  IPITASVEIQEP-----SSRRRVNSLPEVLPILNSNEAKTVQTDVHFLKEGCGDDNVCNS  IPVSLSIAIHTP----EARRRQGRSLPDLLPILNASEPNTVHSEVQFLKEGCGSDNVCHS  IPLSVDVKISGSDSGSQSRRRQGRALPDLVPVLNSNEPET--VKAEFLKEGCGEDNICNS  IPIAVSVNIAGLESGSSST-RKERALPDLIPILNSNESETKITKVEFLKEGCGEDNECHS  IPISVSVKIAGLESP--SK-RKETALPDLTPILNSNESETEITKVEFLKEGCGEDNECHS | 646  644  643  637  634  592  634  634  634  518  633  634  634  634  634  634  631  638  633  542 |
| --- | --- | --- |

| fugu  shortfin_molly  gar  frog  elephant  bear  dog  cow  pig  guinea_pig  chinchilla  human  gorilla  monkey  mouse  rat  python  alligator  chicken  crow | NLKMEYKLHYK---QDPYSPLPVENNIPVFHLSYQRKDLAVQITVSN---------TNGD  NLMIEYRYVYRTTDVDDFTPLNMENGVPVFSFTSQ-KHIALEVTVTN---------PHGD  NLDFKYKFCSREHNRDVFHPLPVENGVPVISLSNQ-EDIALEVTVTNMPSRD----RDGD  NMQLKYKFLTREGSHDSFTELKQENGIPVLALKNQ-KEIALEVTVTNKPSNPANPKLDGD  NLKLEYKFCTREGSQDKFSYLPIQKGVPELVLKDQ-KDIALEITVTNSPSNPRNPTNDGD  NLKLEYKFCTREGNQDKFSYLPIQKGVPELVLKDQ-KDIALEITVTNSPSDPRDPAKDGD  NLKLEYKFCTREGSQDKFSYLPIQKGVPELVLKDQ-KDIALEITVTNSPSNPRDPTKDGD  NLKLEYKFCTREGNQDKFSYLPIHKGVPELVLKDQ-KDIALEITVTNSPSNPKNPTKDGD  NLKLEYKFCTREGNQDKFSYLPIQKGVPELVLKDQ-KDIALEITVTNSPSXPRNPTKDGD  NLKLEYKFCTREGNQDKFSYLPIQKGVPELVLKDQ-KDIALEITVTNIPSDPRNPKKDGD  NLKLEYKFCTREGSQDKFSYLPIQKGVPELVLKDQ-KDIALEITVTNIPSDPRNPKKDGD  NLKLEYKFCTREGNQDKFSYLPIQKGVPELVLKDQ-KDIALEITVTNSPSNPRNPTKDGD  NLKLEYKFCTREGNQDKFSYLPIQKGVPELVLKDQ-KDIALEITVTNSPSNPRNPTKDGD  NLKLEYKFCTREGNQDKFSYLPIQKGVPELVLKDQ-KDIALEITVTNSPSNPRNPTKDGD  NLKLEYKFGTREGNQDKFSYLPIQKGIPELVLKDQ-KDIALEITVTNSPSDPRNPRKDGD  NLKLEYKFGTREGNQDKFSYLPIQKGIPELVLKDQ-KDIALEITVTNGPSDPRNPRKDGD  NLKMQYRFCTKEGNEDRFAYFPMENNVPMLVLKDQ-KDIALEITVTNSPSNAMYPLKDGE  NLKLQYRFCTREGNEDRFSYLPIENSVPVLVLKDQ-KDIALEITVTNNPSDAKNPKKDGE  NLKLQYRFCTREGNEDRFTYLPIENGIPVLVLKDQ-KDIALEITVTNNPSDARNPQKDGE  NLKLQYRFCTREGNEDRFTYLPLENGMPVLVLKDQ-KDIALEITVTNNPSDVKYPQKDGE | 694  694  698  696  693  651  693  693  693  577  692  693  693  693  693  693  690  697  692  601 |
| --- | --- | --- |

| fugu  shortfin_molly  gar  frog  elephant  bear  dog  cow  pig  guinea_pig  chinchilla  human  gorilla  monkey  mouse  rat  python  alligator  chicken  crow | DAYEARLLATFPKMLSYSGVRSHSATTEKPIICTANQDGSQADCELGNPFKRDSKQVTFY  DAYEASVTANFPSSLTYSAYRVS--PEKLQITCIANKNGSMADCELGNPFKRDS-ETTFY  DAYEAKLVALLPDTLSYSAARTLNAPSDKQVTCTANQNGSRADCDLGNPFKSKS-VVTFY  DAHEAQLTAELPSSLSYSKYVELNPQLDKPLICASNPNGSVVFCELGNPFKRNA-NVTFY  DAHEAKLIATFPDTLTYSAYRELRSFPEKQLSCVANQNGSQADCELGNPFKRNS-NVTFY  DAHEAKLVAAFPDTLTYSAYRELRAFPEKQLSCVANQNGSQADCELGNPFKRNS-SVTFY  DAHEAKLVATFPDTLTYSAYRELRAFPEKQLSCVANQNGSQVDCELGNPFKRNS-SVTFY  DAHEAKLIATFPDTLTYSAYRELRAFPEKQLSCVANQNGSQADCELGNPFKRNS-SVTFY  DAHEAKLIATFPDTLTYSAYRELRAFPEKQLSCVANQNGSQADCELGNPFKRNS-SVTFY  DAHEAKLIATFPDTLTYSAYRELRAFPEKQLSCVANQNGSQADCELGNPFKRNS-SVTFY  DAHEAKLIATFPDTLTYSAYRELRAFPEKQLSCVANQNGSQADCELGNPFKRNS-SVTFY  DAHEAKLIATFPDTLTYSAYRELRAFPEKQLSCVANQNGSQADCELGNPFKRNS-NVTFY  DAHEAKLIATFPDTLTYSAYRELRAFPEKQLSCVANQNGSQADCELGNPFKRNS-NVTFY  DAHEAKLIATFPDTLTYSAYRELRAFPEKQLSCVANQNGSQADCELGNPFKRNS-SVTFY  DAHEAKLIATFPDTLTYSAYRELRAFPEKQLSCVANQNGSQADCELGNPFKRNS-SVTFY  DAHEAKLIATFPDTLTYSAYRELRAFPEKQLSCVANQNGSQADCELGNPFKRNS-SVTFY  DAHEARFVATFPDSLTYSAFREWKSFPEKLLTCGANPNGSQVECELGNPFKRNS-TVTFY  DAHEAKLIATFPDSLTYSTFRELRAYPEKQLTCGANQNGSQAECELGNPFKRNS-NVTFY  DAYEAKLIATFPDSLTYSAFREMRGYPEKQLTCGANQNGSQAECELGNPFKRNS-NVTFY  DAYEAKLIATFPDSLTYSAFREMRGYPEKQLTCGANQNGSQAECELGNPFKRNS-NVTFY | 754  751  757  755  752  710  752  752  752  636  751  752  752  752  752  752  749  756  751  660 |
| --- | --- | --- |

| fugu  shortfin_molly  gar  frog  elephant  bear  dog  cow  pig  guinea_pig  chinchilla  human  gorilla  monkey  mouse  rat  python  alligator  chicken  crow | IILSTTNMSVDTTEINIDLQLQTTSVQD-INPVQVKAKVVIEFPLSVSGQARPNQVSFGG  LILGTAGISASTSELEVELVLKTTSNQDNLMPVKAKAKVAIVLQMSLSGQVQPSQVYFSG  IILSTSGITLDTTQVEIDLQLETASEQKNLPKLKANAKVLIALLLSVSGVARPSQVYFGG  LILSTNEISVDTNELDIGLNLKTTSSQANLAPVAAKARVVVELLLSVSGVAKPSQVYFGG  LILSTTEVTFDTTDLDINLKLETTSNQDNLASITAKAKVVIELLLSVSGVAKPSQVYFGG  LILSTTEVTFDTTDLDINLKLETTSNQDNLASITATAKVVIELLLSVSGVAKPSQVYFGG  LILSTTEVTFDTTDLDINLKLETTSNQDNLASITATAKVVIELLLSVSGVAKPSQVYFGG  LILSTTEVTFDTTDLDINLKLETTSNQDNLASITATAKVVIELLLSVSGVAKPSQVYFGG  LILSTTEVTFDTTDLDINLKLETTSNQDNLASITATAKVVIELLLSVSGVAKPSQVYFGG  LILSTTELTFDTTDLDINLKLETTSNQDNLASITAKAKVVIELLLSVSGVAKPSQVYFGG  LILSTTEVTFDTTDLDINLKLETTSNQDNLASITAKAKVVIELLLSVSGVAKPSQVYFGG  LVLSTTEVTFDTPDLDINLKLETTSNQDNLAPITAKAKVVIELLLSVSGVAKPSQVYFGG  LVLSTTEVTFDTPDLDINLKLETTSNQDNLAPITAKAKVVIELLLSVSGVAKPSQVYFGG  LVLSTTEVTFDTPDLDINLKLETTSNQDNLAPITAKAKVVIELLLSVSGVAKPSQVYFGG  LILSTTEVTFDTTDLDINLKLETTSNQDNLAPITAKAKVVIELLLSVSGVAKPSQVYFGG  LILSTTEVTFDTTDLDINLKLETTSNQDNLAPITAKAKVVIELLLSVSGVAKPSQVYFGG  VILSTTKVNVDTKELDINLKLETTSNQTNLEPITAKAKVVIELLLSLTGVAKPSQVYFGG  LVLSTTKVNVDTTDLDITLELETTSNQVNLVPITATAKVVIQLLLSVTGVAKPSQVYFGG  LILSTTKVNVDTTDLDINLKLETTSTQVNSTAITASAKVVLELLLSLTGVAKPSQVYFGG  LILSTTKVNVDTTDLDINLKLETTSTQVNLTPITASAKVVLELLLSLTGVAKPSQVYFGG | 813  811  817  815  812  770  812  812  812  696  811  812  812  812  812  812  809  816  811  720 |
| --- | --- | --- |

| fugu  shortfin_molly  gar  frog  elephant  bear  dog  cow  pig  guinea_pig  chinchilla  human  gorilla  monkey  mouse  rat  python  alligator  chicken  crow | VVKGESAMKTEEEIGSLINFTFTINNLWKS---AVKASLNIHWPKWNKDGKWLLYLVRIS  NVRGEMAMKTESDVGSAVAFQFRIVNLGKRLTELGTATLQIEWPKSTKYEKWLLYLMKIS  TVKGESAMRKEDDIGSLVDYEFRVINLGKPLKSFGSASLNVQWPKEAANHKWLLYLMKVS  NVLGESAMKSEDDIGNLIDFDFRVTNFGRPLKALGTTFLNIQWPKEIYNGKWLLYLVKME  TVVGEQAMKSEDEVGSLIEYEFSVINLGKPLKNLGTATLNIQWPKEISNGKWLLYLVKVE  TVVGEQAMKSEDEVGSLIEYEFRVINLGKPLKNLGTATLNIQWPKEISNGKWLLYLVKVE  TVVGEQAMKSEDEVGSLIEYEFRVINLGKPLKNLGTATLNIQWPKEISNGKWLLYLVKVE  TVVGEQAMKSEEDVGSLIEYEFRVINLGKPLKNLGTATLNIQWPKEISNGKWLLYLVKVE  TVVGEQAMKSEDEVGSLIEYEFRVINLGKPLKNLGTATLNIQWPKEISNGKWLLYLVKVE  TVVGEQAMKSEDEVGSLIEYEFKVINLGKPLKNLGSATLNIQWPKEISNGKWLLYLVKVE  TVVGEQAMKSEDEVGSLIEYEFKVINLGKPLKNLGSATLNIQWPKEISNGKWLLYLVKVE  TVVGEQAMKSEDEVGSLIEYEFRVINLGKPLTNLGTATLNIQWPKEISNGKWLLYLVKVE  TVVGEQAMKSEDEVGSLIEYEFRVINLGKPLTNLGTATLNIQWPKEISNGKWLLYLVKVE  TVVGEQAMKSEDEVGSLIEYEFRVINLGKPLKNLGTATLNIQWPKEISNGKWLLYLVKVE  TVVGEQAMKSEDEVGSLIEYEFRVINLGKPLKNLGTATLNIQWPKEISNGKWLLYLMKVE  TVVGEQAMKSEDEVGSLIEYEFRVINLGKPLKNLGTATLNIQWPKEISNGKWLLYLMKVE  NIIGESAMKSEDDIGSLIEYEFRITNLGKPLKAFGTAFLEIMWPKELRGGKWLLYLVSIH  NIIGESAMKSEDDIGNLIEYEFRITNLGRPLKTFGTASLDIQWPKEIDNGKWLLYLMKVE  NIVGESAMKSEDNIGNLIEYEFRVTNLGRPLKTFGTASLDIQWPKEISNGKWLLYLMKIE  NIIGESAMKSEDDIGNLIEYEFRVTNLGRPLKTFGTASLDIQWPKEISNGKWLLYLMKID | 870  871  877  875  872  830  872  872  872  756  871  872  872  872  872  872  869  876  871  780 |
| --- | --- | --- |

| fugu  shortfin_molly  gar  frog  elephant  bear  dog  cow  pig  guinea_pig  chinchilla  human  gorilla  monkey  mouse  rat  python  alligator  chicken  crow | STGPQEILCSPLSEVNSLKLQQDLASPRSKREIGERKRVGK----ATLLSGKNKVLSCDK  STGVERIHCTPNEE-NLLNLEPVK-TRK--RRAAEITGERTDYAFSRL-DHNKETLSCDG  ATGTKRIDCTPRGEVNSLNLTPSS-SSRKKRETEQRQTPGEGKFSLFMDKRKYTTLSCGN  SEGLDKMECQPAGEINKLKLLESG-KTRHRREIGESQSGTSDKSFSLFSERKYMTLDCNN  SKGLEKIGCEPPSEINFLKLKESH-NSRKKREIAEKQIDD-RRKFSLFAERKYQTLNCSV  SKGLEKLDCEPRSEINFLNLKEFH-NSRRKREIAEKQTDD-SRKFSLFAERKYQTLNCST  SKGLEKIACEPRGEINFLKLKESH-NSRRKREIAEKQIED-SRKFSLFAERKYQTLNCST  SKGLEKITCEPQGEINFLKLTEAH-NSRRKREIAERQTDD-GRKFSLFSERKYQTLNCSV  SKGLDKVACQPQREINSLQLKESH-NSRRKREIAERQRDD-GRKFSLFAERKYQTLNCSV  SKGLEKIVCEPPDEINPLKLKESH-KSRMKREIAEKQIDD-RRKFSLFTERKYQTLNCSV  SKGLEKIVCEPQNEINVLNLKESH-KSRMKREIAEKQIDD-RRKFSLFTERKYQTLNCSV  SKGLEKVTCEPQKEINSLNLTESH-NSRKKREITEKQIDD-NRKFSLFAERKYQTLNCSV  SKGLEKVTCEPQKEINSLNLTESH-NSRKKREITEKQIDD-NRKFSLFAERKYQTLNCSV  SKGLEKITCEPQKEINFLNLKESH-NSRKKREITEKQIDD-SRKFSLFAERKYQTLNCSV  SKGLEQIVCEPHNEINYLKLKESH-NSRKKRELPEKQIDD-SRKFSLFPERKYQTLNCSV  SKGLEQVVCEPHNEINFLKLKESH-NSRKKRELPEKQIDD-SRKFSLFSERKYQTLNCSV  SKELGELQCAPQNEINPLKVQVSK-NSRTKREVAEKQTEG-SKGFSLFTERKYKTLECNE  SKELGIIPCEPRNEINYLQLRDSH-NSRRKREIAEKQIND-GKKFSLFSERKYKTLDCRG  SKGLEKVSCQPQNEINVLHVAESH-NSRRKREIAEKQLTD-SKTFSLFSERKYKTLDCKV  SKGLEKVSCQPENEINSLHVAETH-NSRRKREVAEKQITD-SKAFSLFSERKYKTLNCNV | 926  926  936  934  930  888  930  930  930  814  929  930  930  930  930  930  927  934  929  838 |
| --- | --- | --- |

| fugu  shortfin_molly  gar  frog  elephant  bear  dog  cow  pig  guinea_pig  chinchilla  human  gorilla  monkey  mouse  rat  python  alligator  chicken  crow | GSRCVVLECPLQGVDGT-TVELRSRLWNSTFIEDYAALLHLDIVVRASLVLHSQAKNIIL  TAKCVTIRCALGGLDRNALITLNSRLWNSTFIEDYPKFHHVEVVVKASLHVGSATTNTML  GTQCVEIRCPLQGLDNNPIIVLRSRLWNSTFLEDYSTLNYLDILVKASLSLDASAKNIVL  QAKCVTIKCPLHGMDSNAIIKARSRLWNSTFLEEYSKMNYLDILVKVFISVDTAAQNIKL  NVKCVNIKCPLQGLDSKASVILRSRLWNSTFLEEYSKMNYLDILTRASIDVTAAAENIKL  NVNCVNISCPLRGLDSKASVILRSRLWNSTFLEEYSKMNYLDILMRASIDVAAAAENIKL  NVRCVNISCPLRGLDSKASVVLRSRLWNSTFLEEYSKMNYLDILVRASIDVAATAENIKL  NVNCVNIKCPLRGLDSKASVVLRSRLWNSTFLEEYSKMNYLDILVRASIDVIAATENIKL  NVNCVNIRCPLQGLDSKASVVLRSRLWNSTFLEEYSKMNYLDILVRASIDMTAATENIKL  NVKCVNIRCPLQGLDSKAALVLRSRLWNGTFLEEYSKMNYLDILVRAYIEVTAAAGNIKL  NVNCVNIRCPLRGLDSKAALVLRSRLWNGTFLEEYSKMNYLDILVRASIEVTAAAGNIKL  NVNCVNIRCPLRGLDSKASLILRSRLWNSTFLEEYSKLNYLDILMRAFIDVTAAAENIRL  NVNCVNIRCPLRGLDSKASLILRSRLWNSTFLEEYSKLNYLDILMRAFIDVTAAAENIRL  NVNCVNIRCPLRGLDSKASLILRSRLWNSTFLEEYSKLNYLDILVRAFIDVTAAAENIKL  NVRCVNIRCPLRGLDSKASLVLRSRLWNSTFLEEYSKLNYLDIL**L**RASIDVTAAAQNIKL  NVRCVNIRCPLRGLDSKASLVLRSRLWNSTFLEEYSKLNYLDILVRASIDVTAAAQNIKL  EANCVNITCSLQGLDSKAIVVLRSRLWNSTFLEEFYKMNYLDIPVRARIGISA-ADNVKL  SARCVNIKCPLQGLDSKALVVLRSRLWNSTFLEEYSKLNYLDIVVRATISIPAAAENVKL  NAQCVDIRCPLKGFDSKASILLRSRLWNSTFLEEFSKMNYLDILVRASISVPAAAKNVKL  NARCVDIKCPLKGFDSKASIVLRSRLWNSTFLEEFSTMNYLDILVRASISVPAAAKNVKL | 985  986  996  994  990  948  990  990  990  874  989  990  990  990  990  990  986  994  989  898 |
| --- | --- | --- |

| fugu  shortfin_molly  gar  frog  elephant  bear  dog  cow  pig  guinea_pig  chinchilla  human  gorilla  monkey  mouse  rat  python  alligator  chicken  crow | RTPDTEVMLTVSPEPTVAQHTGVPWWIILVAVLAGILILALLVFLLWKCGFFTRPLQDDS  KNPETTVKLTVFPERRSAQYGGVPWWIIVLSILLGLLLLALLAFLLWKCGFFNRAKYEDK  KNAETQVKLTVFPEKTVAKYTGAPWWIILVAILAGILMLALLVFLLWKCGFFKRSKYDDS  ANEGYQVRVTVFPEKTVAQYSGVPWWIIFVAILAGILMLALLVFLLWKCGFFKRSRYDDS  PNAGTQVRVTVFPSKTAAQYSGVSWWIILVAILAGILMLGLLVFLLWKCGFFKRSRYDDS  PNAGTQIRVTVFPSKTVAQYSGIPWWIILVAILAGILMLALLVFLLWKCGFFKRSRYDDS  PNAGTQIRVTVFPSKTVAQYSGVPWWIILVAILAGILMLALLVFILWKCGFFKRSRYDDS  PNAGTQVRVTVFPSKTVAQHSGVPWWIILVAILAGILMLGLLVFLLWKCGFFKRSRYDDS  PNAGTQVRVTVFPSKTAAQYSGVPWWIILVAILAGILMLALLVFLLWKCGFFKRSRYDDS  PHAGTQVRVTVFPSKTVAQYSGIPWWVILVSILAGILMLALLVIVLWKCGFFNRARYDDS  PHAGTQVRVTVFPSKTVAQYSGIPWWIILVAILAGILMLALLVFLLWKCGFFKRSRYDDS  PNAGTQVRVTVFPSKTVAQYSGVPWWIILVAILAGILMLALLVFILWKCGFFKRSRYDDS  PNAGTQVRVTVFPSKTVAQYSGVPWWIILVAILAGILMLALLVFILWKCGFFKRSRYDDS  PNAGTQVRVTVFPSKTVAQYSGVPWWIILVAILAGILMLALLVFILWKCGFFKRSRYDDS  PHAGTQVRVTVFPSKTVAQYSGVAWWIILLAVLAGILMLALLVFLLWKCGFFKRSRYDDS  PHAGTQVRVTVFPSKTVAQYSGIAWWIILLAVLAGILMLALLVFLLWKCGFFKRSRYDDS  TNEAANVRVTVFPAKTVALYKGVPWWIILVAILAGLLMLALLVFLLWKCGFFQRSRYDDS  TNETAQVRVTVFPAKTVAHYTGVPWWIILVAILAGILMLALLVFLLWKCGFFQRSRYDDS  TNEAAQVRVTVFPAKPVALYTGVPWWIIAVAIFAGVLMLALLVFLLWKCGFFQRSRYDDS  TNEVTQVRVTVFPAKPVALYTGVPWWIIAVAILAGILMLALLVFLLWKCGFFRRSRYEDS | 1045  1046  1056  1054  1050  1008  1050  1050  1050  934  1049  1050  1050  1050  1050  1050  1046  1054  1049  958 |
| --- | --- | --- |

| fugu  shortfin_molly  gar  frog  elephant  bear  dog  cow  pig  guinea_pig  chinchilla  human  gorilla  monkey  mouse  rat  python  alligator  chicken  crow | VPRYHAVRIKKETPEYKDGFKR-DSFEKKPWVTTWSDHESYS  VPSYSAVRIRREERAVHSAKDNWGNLETKPWMTTWHDKEHYS  VPRYHAVRIRKEERQFKDGKTKLVKLEKKQWMTTWNENESYS  VPRYHAVRISKEEREYKDGKLP-KNSEKKQWVTKWNENESYS  VPRYHAVRIRKEEREIQDEKYNNDNLEKKQWIAKWTENESYS  VPX--------EEREIKDEKYN-DNLEKKQWITRWNENESYS  VPRYHAVRIRKEEREIKDEKYN-DNLEKKQWITSWNENESYS  VPRYHAVRIRKEEREIPDEKYN-DNPEKKQWITKWNENESYS  VPRYHAVRIRKEEREIKDEKYN-DNLEKKQWITRWNENESYS  VPRYYAVRIRKEEREIKDEKYN-DNLEKKQWITKWNENESYS  VPRYYAVRIRKEEREIKDEKYN-DNLEKKQWITKWNENESYS  VPRYHAVRIRKEEREIKDEKYI-DNLEKKQWITKWNENESYS  VPRYHAVRIRKEEREIKDEKYI-DNLEKKQWITKWNENESYS  VPRYHAVRIRKEEREIKDEKYI-DNLEKKQWITKWNENESYS  IPRYHAVRIRKEEREIKDEKHM-DNLEKKQWITKWNENESYS  VPRYHAVRIRKEEREINDEKHT-DNPE-KKWPPKWNENESYS  IPRYHAVRIRKEERQIKDGKSK-ENHAKKQWITKWNGNESYS  VPRYHAVRIRKEERQIKDGKYK-DDHEKKQWITKWSENESYS  VPRYHAVRIRKEEREIKDGKCK-D-LETKQWFTKWNENESYS  VPCYHAVRIRKEERHIKDGNCK-D-LETKQWFTKWNENESYS | 1086  1088  1098  1095  1092  1041  1091  1091  1091  975  1090  1091  1091  1091  1091  1090  1087  1095  1089  998 |
| --- | --- | --- |
