## Supplemental Figure 3 for "Analysis of a genetic region affecting mouse body weight"

| Frog  Mouse  Rat  guinea_pig  chinchilla  cow  bear  dog  human  gorilla  snubnosed_monkey  pig  elephant  python  alligator  crow  chicken  grouse  spotted_gar  shortfin_molly  fugu | ---------------------------------MRL---------------LRAL-----  ---------------------------------MRLARLLRGGTSVRPLCA----VPCAS  ---------------------------------MRLARLLRGGTSVRPLCA----VPCAS  ---------------------------------MKLTRLLRGAASARPGPGPRATHPSAS  ---------------------------------MKLTRLLRGAASARPRPGPRAAHPSAS  ---------------------------------MRLARLLRVATSAGPGPELRAAGPSVS  ------------------------------------------------------------  ------------------------------------------------------------  ---------------------------------MRLARLLRGAALAGPGPGLRAAG--FS  MPAPAARHVPHVPLGRGAGKPGALAVASLAGLGMRLARLLRGAALAGPGPGLRAAG--SS  ---------------------------------MRLARLLRGAALAVPGPGLRAAG--PS  ---------------------------------MRLARLLRGAASAGPGPGLRAACPSAS  ------------------------------------------------------------  ---------------------------------MRL---------------LRALL----  ------------------------------------------------------------  ------------------------------------------------------------  ---------------------------------MRL---------------LRALL----  ------------------------------------------------------------  ---------------------------------MRL------------------------  ---------------------------------MRL------------------------  ---------------------------------MRI------------------------ | 7  23  23  27  27  27  0  0  25  58  25  27  0  8  0  0  8  0  3  3  3 |
| --- | --- | --- |

| Frog  Mouse  Rat  guinea_pig  chinchilla  cow  bear  dog  human  gorilla  snubnosed_monkey  pig  elephant  python  alligator  crow  chicken  grouse  spotted_gar  shortfin_molly  fugu | --------MRSAPLSSRNTPSLVDFYSKFSPSP----------LSMKQFLDFG-----SV  RSL**AS**ASASGSGPASELGVPGQVDFYARFSPSP----------LSMKQFLDFG-----SV  RSLASDSASGSGPASESGVPGQVDFYARFSPSP----------LSMKQFLDFG-----SV  RS--LASDSGTGPASDRSVPGQVDFYARFSPSP----------LSMKQFLDFG-----SV  RS--LTSDSGSGPAPDRSLPGQVDFYARFSPSP----------LSMKQFLDFG-----SG  RS--LTSDSGSGPAPERGVPGQVDFYARFS---PSP----LSM---KQFLDFG-----SV  -----------------------------------------------------------M  ----MRREA--------G----YNWPERFTECLPCHLLNSVSIRGTFVSSSAR-----SM  RS--FSSDSGSSPASERGVPGQVDFYARFSPS---P-------LSMKQFLDFG-----SV  RS--FSSDSGSSPASERGVPGQVDFYARFSPS---P-------LSMKQFLDFG-----SV  RT--FSSDSGSGPASERGVPGQVDFYARFSPS---P-------LSMKQFLDFG-----SV  RS--LTSDSGSGPAPELGVPGQVDFYARFSPS---P-------LSMKQFLDFG-----SV  ---------------------------------------------MKQFLDFG-----SV  ----------RSVSPAATIPQQVDFYCRFSPSP----------LSMKQFLDFG-----SD  ------------------------------------------------------------  ------------------------------------------------------------  ----------RCASP-GSIPQQVDFYSRFSPSP----------LSMKQFLDFG-----SE  ------------------------------------------------------------  ---------LTFLLKNGSLGKQVDFYSRFSPSP----------LSMKQFLDFG-----SE  ---------LRFLRSSVSIGKDIDFYSRFSPSP----------LSMKQFLDFGKSCTGSE  ---------LRFLRSSISAGKDIDYYSKFSPSP----------LSMKQFLDFG-----SE | 44  68  68  70  70  70  1  39  68  101  68  70  10  43  0  0  42  0  39  44  39 |
| --- | --- | --- |

| Frog  Mouse  Rat  guinea_pig  chinchilla  cow  bear  dog  human  gorilla  snubnosed_monkey  pig  elephant  python  alligator  crow  chicken  grouse  spotted_gar  shortfin_molly  fugu | NACEKTSFIFLRHELPVRLANIMKEINLLPDNLLKMPSIKLVQSWYVQSFQEIIDFKDTN  NACEKTSFMFLRQELPVRLANIMKEISLLPDNLLRTPSVQLVQSWYIQSLQELLDFKDKS  NACEKTSFMFLRQELPVRLANIMKEISLLPDNLLRTPSVQLVQSWYIQSLQELLDFKDKS  NACEKTSFMFLRQELPVRLANIMKEISLLPDNLLRTPSVQLVQSWYIQSLQELLDFKDKS  NACEKTSFMFLRQELPVRLANIMKEISLLPDNLLRTPSVQLVQSWYIQSLQELLDFKDKS  NACEKTSFMFLRQELPVRLANIMKEISLLPDNLLRTPSVQLVQSWYIQSLQELLEFKDKS  NACEKTSFMFLRQELPVRLANIMKEISLLPDNLLRTPSVQLVQSWYIQSLQELLEFKDKS  NACEKTSFMFLRQELPVRLANIMKEISLLPDNLLRTPSVQLVQSWYIQSLQELLEFKDKS  NACEKTSFMFLRQELPVRLANIMKEISLLPDNLLRTPSVQLVQSWYIQSLQELLDFKDKS  NACEKTSFMFLRQELPVRLANIMKEISLLPDNLLRTPSVQLVQSWYIQSLQELLDFKDKS  NACEKTSFMFLRQELPVRLANIMKEISLLPDNLLRTPSVQLVQSWYIQSLQELLDFKDKS  NACEKTSFMFLRQELPVRLANIMKEISLLPDNLLRTPSVQLVQSWYIQSLQELLDFKDKS  DACEKTSFMFLRQELPVRLANIMKEISLLPDNLLRTPSVQLVQSWYIQSLQELLDFKDKS  NACEKTSFMFLRQELPVRLANIMKEISLLPDNLLKTPSVQLVQSWYVQSLQEILDFKDKN  --------MFLRQELPVRLANIMKEISLLPDNLLRTPSVQLVQSWYDQSLQEILDFKDKN  --------MFLRQELPVRLANIMKEISLLPDNLLRTPSVQLVQSWYVQSLQEILDFKDKS  NACEKTSFMFLRQELPVRLANIMKEISLLPDNLLRTPSVQLVQSWYVQSLQEILDFKDKS  --------MFLRQELPVRLANIMKEISLLPDNLLRTPSVQLVQSWYVQSLQEILDFKDKS  NACERTSFVFLRQELPVRLANIMKEINLLPDNLLKTPSVQLVQSWYIQSFQEILDFKERK  NACEKTSFAFLRQELPVRLANIMKEINLLPDNLLRTPSVQLVQSWYMQSFQDILDFKDKN  NACEKTSFAFLRQELPVRLANIMKEINLLPDNLLRTPSVRLVQSWYMQSFQDILEFRDRN | 104  128  128  130  130  130  61  99  128  161  128  130  70  103  52  52  102  52  99  104  99 |
| --- | --- | --- |

| Frog  Mouse  Rat  guinea_pig  chinchilla  cow  bear  dog  human  gorilla  snubnosed_monkey  pig  elephant  python  alligator  crow  chicken  grouse  spotted_gar  shortfin_molly  fugu | AEDLNTVQKFTDTVITIRNRHNDVIPTMAQGVVEFKDSFGVDPVTSQNVQYFLDRFYMSR  AEDAKTIYEFTDTVIRIRNRHNDVIPTMAQGVTEYKESFGVDPVTSQNVQYFLDRFYMSR  AEDAKTIYEFTDTVIRIRNRHNDVIPTMAQGVTEYKESFGVDPVTSQNVQYFLDRFYMSR  AEDAKAIYDFTDTVIRIRNRHNDVIPTMAQGVIEYKESFGVDPVTSQNVQYFLDRFYMSR  AEDAKTIYDFTDTVIRIRNRHNDVIPTMAQGVIEYKESFGVDPVTIQNVQYFLDRFYMSR  AEDAKTIYDFTDTVIRIRNRHNDVIPTMAEGVVEYKESFGVDPVTSQNVQYFLDRFYMSR  AEDAKTIYDFTDTVIRIRNRHNDVIPTMAQGVIEYKESFGVDPVTSQNVQYFLDRFYMSR  AEEAKTIYDFTDTVIRIRNRHNDVIPTMAQGVIEYKESFGVDPVTSQNVQYFLDRFYMSR  AEDAKAIYDFTDTVIRIRNRHNDVIPTMAQGVIEYKESFGVDPVTSQNVQYFLDRFYMSR  AEDAKAIYDFTDTVIRIRNRHNDVIPTMAQGVIEYKESFGVDPVTSQNVQYFLDRFYMSR  AEDAKAIYDFTDTVIRIRNRHNDVIPTMAQGVIEYKESFGVDPVTSQNVQYFLDRFYMSR  AEDAKTIYDFTDTVIRIRNRHNDVIPTMAQGVVEYKESFGVDPVTSQNVQYFLDRFYMSR  AEDAKTIYDFTDTVIRIRNRHNDVIPTMAQGVVEYKESFGVDPVTSQNVQYFLDRFYMSR  AEDTEAVSRFTDTVITIRNRHNDVIPTMAQGVIEYKDNYGVDPVTSQNVQYFLDRFFMSR  AEDSETIHSFTDTVIKIRNRHNDVIPTMAQGVVEYKESFGIDPVTSQNVQYFLDRFYMSR  SEDSGAVHSFTDTVIKIRNRHNDVIPTMAQGVIEYKESFGIDPVTSQNVQYFLDRFYMSR  SEDSGAIHSFTDTVIKIRNRHNDVIPTMAQGVIEYKESFGIDPVTSQNVQYFLDRFYMSR  SEDSGTIHSFTDTVIKIRNRHNDVIPTMAQGVIEYKESFGIDPVTSQNVQYFLDRFYMSR  AEDDKVIYDFTDAVIKIRNRHNDVIPTLAQGVVEYKESYGIDPVTSQNVQYFLDRFFMSR  ADDEKVTCDFTDAVIKIRNRHNDVIPTMAQGLVEYKETYGTDPVVSQNVQYFLDRFYMSR  AEDEKVTHDFTNAVIKIRNRHNDVIPTMAQGVVEYKETYGTDPVVSQNVQYFLDRFYMSR | 164  188  188  190  190  190  121  159  188  221  188  190  130  163  112  112  162  112  159  164  159 |
| --- | --- | --- |

| Frog  Mouse  Rat  guinea_pig  chinchilla  cow  bear  dog  human  gorilla  snubnosed_monkey  pig  elephant  python  alligator  crow  chicken  grouse  spotted_gar  shortfin_molly  fugu | ISIRMLLNQHTLLFGGEVKVNPAHPKHIGSIDPACNVVDVVKDGYENAKHLCDLYYMSSP  ISIRMLLNQHSLLFGGK--GSPSHRKHIGSINPNCDVVEVIKDGYENARRLCDLYYVNSP  ISIRMLLNQHSLLFGGK--GSPSHRKHIGSINPNCDVVEVIKDGYENARRLCDLYYINSP  ISIRMLLNQHSLLFGGK--GSPSHRKHIGSINPNCDVVEVIKDGYENARRLCDLYYINSP  ISIRMLLNQHSLLFGGK--GSPSHRKHIGSINPNCDVVEVIKDGYENARRLCDLYYINSP  ISIRMLLNQHSLLFGGKGKGSLSHRKHVGSINPNCSVVEVIKDGYENARRLCDLYYINSP  ISIRMLLNQHSLLFGGKGKGSPAHRKHIGSINPNCNVVEVIKDGYENARRLCDLYYINSP  ISIRMLLNQHSLLFGGKGKGSPAHRKHIGSINPNCDVVEVIKDGYENARRLCDLYYINSP  ISIRMLLNQHSLLFGGKGKGSPSHRKHIGSINPNCNVLEVIKDGYENARRLCDLYYINSP  ISIRMLLNQHSLLFGGKGKGSPSHRKHIGSINPNCNVVEVIKDGYENARRLCDLYYINSP  ISIRMLLNQHSLLFGGKGKGSPSHRKHIGSINPNCNVVEVIKDGYENARRLCDLYYINSP  ISIRMLLNQHSLLFGGKGKGSLSHQKHIGSINPNCNVVEVIKDGYENARRLCDLYYINSP  ISIRMLLNQHSLLFGGKGKGSPSHRKHIGSINPHCNVVEVIKDGYENARRLCDLYYINSP  ISIRMLLNQHTLLFGGKVEVNPAHPKHIGSIDPKCNVVEVIKDGYENAKSLCDLYYMSSP  ISIRMLLNQHSLLFGGKIKVNPAHPKHIGSINPNCDVVGVIKDGYENAKTLCDLYYMSSP  ISIRMLLNQHSLLFGGKI--NPAHPKHIGSIDPNCNVVEVIRDGYENAKRLCDLYYMSSP  ISIRMLLNQHSLLFGGKI--NPAHPKHIGSIDPSCNVVGVIRDGYESAKSLCDLYYMSSP  ISIRMLLNQHSLLFGGKI--NPAHPKHIGSIDPSCNVVGVIRDGYESAKTLCDLYYMSSP  ISIRMLLNQHTLLFGGKVKVNPAHPKQIGSIDPHCRVEEVIKDAYENARRLCDLYYMNSP  ISIRMLLNQHTLLFGGKVKVNPAHPNQIGSIDPHCGVSEVIRDAFENARNLCDRFYMNSP  ISIRMLLNQHTLIFGGK--VNPAHPKQIGGIDPHCRVSDVVRDAFENARNLCDRYYMNSP | 224  246  246  248  248  250  181  219  248  281  248  250  190  223  172  170  220  170  219  224  217 |
| --- | --- | --- |

| Frog  Mouse  Rat  guinea_pig  chinchilla  cow  bear  dog  human  gorilla  snubnosed_monkey  pig  elephant  python  alligator  crow  chicken  grouse  spotted_gar  shortfin_molly  fugu | ELELTEFNAKSPGQPIQVVYVPSHLYHMVFELFKNAMRATMEFQADKGVYPPIKVHVVLG  ELELEELNAKSPGQTIQVVYVPSHLYHMVFELFKNAMRATMEHHADKGVYPPIQVHVTLG  ELELEELNAKSPGQPIQVVYVPSHLYHMVFELFKNAMRATMEHHADKGVYPPIQVHVTLG  ELELEELNAKSPGQPIQVVYVPSHLYHMVFELFKNAMRATMEHHADKGVYPPIQVHVTLG  ELELEELNAKSPGQPIQVVYVPSHLYHMVFELFKNAMRATMEHHADKGVYPPIQVHVTLG  ELELEELNAKSPGQPIQVVYVPSHLYHMVFELFKNAMRATMEHHADKGVYPPIQVHVTLG  ELELEELNAKSPGQPIQVVYVPSHLYHMVFELFKNAMRATMEHHADKGVYPPIQVHITLG  ELELEELNAKSPGQPIQVVYVPSHLYHMVFELFKNAMRATMEHHADKGVYPPIQVHITLG  ELELEELNAKSPGQPIQVVYVPSHLYHMVFELFKNAMRATMEHHANRGVYPPIQVHVTLG  ELELEELNAKSPGQPIQVVYVPSHLYHMVFELFKNAMRATMEHHANRGVYPPIQVHVTLG  ELELGELNAKSPGQPIQVVYVPSHLYHMVFELFKNAMRATMEHHANRGVYPPIQVHVTLG  ELELEELNAKSPGQPIQVVYVPSHLYHMVFELFKNAMRATMEHHADKGVYPPIQVHVTLG  ELELGELNAKSPGQPIQVVYVPSHLYHMVFELFKNAMRATMEYHADKGVYPPIQVLVTLG  ELVLEEQNVKSPGQPMQVVYVPSHLYYMVFELFKNAMRATMEHHADRGIYPPVQVQVTLG  ELVLEEMNVKSPGQPIQVVYVPSHLYHMVFELFKNAMRATMEHNADRCIYPPVHVHVTLG  ELILEELNAKSPGQPMQVVYVPSHLYHMVFELFKNAMRATMEHHADQCIYPAIHVHITLG  ELVLEELNIKSPGQPMQVVYVPSHLYHMVFELFKNAMRATMEHNADRCIYPPIHVHVTLG  ELILKELNVKSPGQPMQVVYVPSHLYHMVFELFKNAMRATMEHNADRCIYPPIHVHVTLG  ELKLEEFNAKERGKPTTVVYVPSHLYHMVFELFKNAMRATMEYHGVIAEYPPVKVQVALG  ELILEEFNAKDEGKPVTVVYVPSHLYHMVFELFKNAMRATMELYGDAMEYPPVHAQVALG  ELVLEEFNVEEKEKPITVVYVPSHLYHMVFELFKNAMRATMELYGDAMEYPAVHAQVALG | 284  306  306  308  308  310  241  279  308  310  250  283  232  230  280  230  280  230  279  284  277 |
| --- | --- | --- |

| Frog  Mouse  Rat  guinea_pig  chinchilla  cow  bear  dog  human  gorilla  snubnosed_monkey  pig  elephant  python  alligator  crow  chicken  grouse  spotted_gar  shortfin_molly  fugu | SEDLTVKLSDRGGGVPLRKIERLFNYMYSTAPLPRMETSRATPLAGFGYGLPISRLYAKY  EEDLTVKMSDRGGGVPLRKIDRLFNYMYSTAPRPRVETSRAVPLAGFGYGLPISRLYAQY  EEDLTVKMSDRGGGVPLRKIDRLFNYMYSTAPRPRVETSRAVPLAGFGYGLPISRLYAQY  NEDLTVKMSDRGGGVPLRKIDRLFNYMYSTAPRPRVETSRAAPLAGFGYGLPISRLYAQY  NEDLTVKMSDRGGGVPLRKIDRLFNYMYSTAPRPRVETSRAAPLAGFGYGLPISRLYAQY  KEDLTVKMSDRGGGVPLRKIDRLFNYMYSTAPRPRVETSRAVPLAGFGYGLPISRLYAQY  NEDLTVKMSDRGGGVPLRKIDRLFNYMYSTAPRPRVETSRAVPLAGFGYGLPISRLYAQY  NEDLTVKMSDRGGGVPLRKIDRLFNYMYSTAPRPRVETSRAVPLAGFGYGLPISRLYAQY  NEDLTVKMSDRGGGVPLRKIDRLFNYMYSTAPRPRVETSRAVPLAGFGYGLPISRLYAQY  NEDLTVKMSDRGGGVPLRKIDRLFNYMYSTAPRPRVETSRAVPLAGFGYGLPISRLYAQY  NEDLTVKMSDRGGGVPLRKIDRLFNYMYSTAPRPRVETSRAVPLAGFGYGLPISRLYAQY  NEDLTVKMSDRGGGVPLRKIDRLFNYMYSTAPRPRVETSRAVPLAGFGYGLPISRLYAQY  NEDLTVKMSDRGGGVPLRKIDRLFNYMYSTAPRPRVETSRAVPLAGFGYGLPISRLYAQY  TEDLTVKMCDRGGGVPLRKIDRLFNYMYSTAPRPRVETSRATPLAGFGYGLPISRLYAQY  NEDLTVKMSDRGGGVPVRKIDRLFNYMYSTAPHPRVETSRATPLAGFGYGLPISRLYAQY  NEDLTVKMSDRGGGVPMRKIDRLFNYMYSTAPRPRVETSRATPLAGFGYGLPISRLYAQY  NEDLTVKMSDRGGGVPMRKIDRLFNYMYSTAPRPRVETSRATPLAGFGYGLPISRLYAQY  NEDLTVKMSDRGGGVPMRKIDRLFNYMYSTAPRPRVETSRATPLAGFGYGLPISRLYAQY  HEDLTVKMSDRGGGVPLRKIERLFSYTYSTAPLPRMETSRATPLAGYGYGLPISRLYARY  SEDLTVKVSDRGGGVPLRKIDRLFTYTYSTAPRPSIDGSRAAPLAGYGYGLPISRLYARY  NEDLTVKVSDRGGGVPLRKIDRLFTYTYSTAPRPSLDGSRAAPLAGYGYGLPISRLYARY | 344  366  366  368  368  370  301  339  368  401  368  370  310  343  292  290  340  290  339  344  337 |
| --- | --- | --- |

| Frog  Mouse  Rat  guinea_pig  chinchilla  cow  bear  dog  human  gorilla  snubnosed_monkey  pig  elephant  python  alligator  crow  chicken  grouse  spotted_gar  shortfin_molly  fugu | FQGDLKLYSLEGYGTDAVIYFKALSTESVERLPVYNKSAWKHYKTNHEADDWCVPSSEPK  FQGDLKLYSLEGYGTDAVIYIKALSTESVERLPVYNKAAWKHYKANHEADDWCVPSREPK  FQGDLKLYSLEGYGTDAVIYIKALSTESIERLPVYNKAAWKHYRTNHEADDWCVPSREPK  FQGDLKLYSLEGYGTDAVIYIKALSTESIERLPVYNKAAWKHYNTNHEADDWCVPSREPK  FQGDLKLYSLEGYGTDAVIYIKALSTESIERLPVYNKAAWKHYNTNHEADDWCVPSREPK  FQGDLKLYSLEGYGTDAVIYIKALSTESIERLPVYNKAAWKHYNTNHEADDWCVPSREPK  FQGDLKLYSLEGYGTDAVIYIKALSTESIERLPVYNKAAWKHYNTNHEADDWCVPSREPK  FQGDLKLYSLEGYGTDAVIYIKALSTESIERLPVYNKAAWKHYNTNHEADDWCVPSREPK  FQGDLKLYSLEGYGTDAVIYIKALSTDSIERLPVYNKAAWKHYNTNHEADDWCVPSREPK  FQGDLKLYSLEGYGTDAVIYIKALSTDSVERLPVYNKAAWKHYNTNHEADDWCVPSREPK  FQGDLKLYSLEGYGTDAVIYIKALSTDSIERLPVYNKAAWKHYNTNHEADDWCVPSREPK  FQGDLKLYSLEGYGTDAVIYIKALSTDSIERLPVYNKAAWKHYNTNHEADDWCVPSREPK  FQGDLKLYSLEGYGTDAVIYIKALSTESIERLPVYNKAAWKHYNTNHEADDWCVPSREPK  FQGDLKLYSLEGYGTDAVIFIKALSTESIEKLPVYNKSAWKHYTANHEADDWCVPSSEPK  FQGDLKLYSFEGYGTDAVIYIKALSTESIERLPVYNKSAWKHYKANHEADDWCVPSSEPK  FQGDLKLYSLEGYGTDAVIYIKALSTESIERLPVYNKAAWKHYKANHEADDWCVPSSEPK  FQGDLKLYSLEGYGTDAVIYIKALSTESIERLPVYNKAAWKHYKANHEADDWCVPSSEPK  FQGDLKLYSLEGYGTDAVIYIKALSTESIERLPVYNKAAWKHYKANHEADDWCVPSSEPK  FQGDLKLYSLEGYGTDAVIYIKALSTDSIERLPVYNRSAWRHYKTIHEADDWCVPSKEPK  FQGDLKLYSLEGYGTDAVIYIRALSTESIERLPVYNKSAWKHYKTMHEADDWCVPSKEPK  FQGDLKLYSLEGHGTDAVIYIRALSTESIERLPVYNKSAWKHYKTIHEADDWCVPSKEPK | 404  426  426  428  428  430  361  399  428  461  428  430  370  403  352  350  400  350  399  404  397 |
| --- | --- | --- |

| Frog  Mouse  Rat  guinea_pig  chinchilla  cow  bear  dog  human  gorilla  snubnosed_monkey  pig  elephant  python  alligator  crow  chicken  grouse  spotted_gar  shortfin_molly  fugu | DMTTFRSN----------------------  DMTTFRSS----------------------  DMTTFRSS----------------------  DMTTFRSA----------------------  DMTTFRST----------------------  DMTTFRSA----------------------  DMTTFRSA----------------------  DMTTFRSA----------------------  DMTTFRSA----------------------  DMTTFRSA----------------------  DMTTFRSA----------------------  DMTTFRSA----------------------  DMTTFRSA----------------------  DMTTFRST----------------------  DMTTFRTWIRLYLNSGNYNSKLDFCKEEKA  DMTTFRSM----------------------  DMTTFRST----------------------  DMTTFRST----------------------  DMTTFRSS----------------------  DLTTFRSF----------------------  DMTTFRSF---------------------- | 412  434  434  436  436  438  369  407  436  469  436  438  378  411  382  358  408  358  407  412  405 |
| --- | --- | --- |
