## Supplementary figures and images for "Analysis of a genetic region affecting mouse body weight"

### Supplemental Figures

#### Slide 1
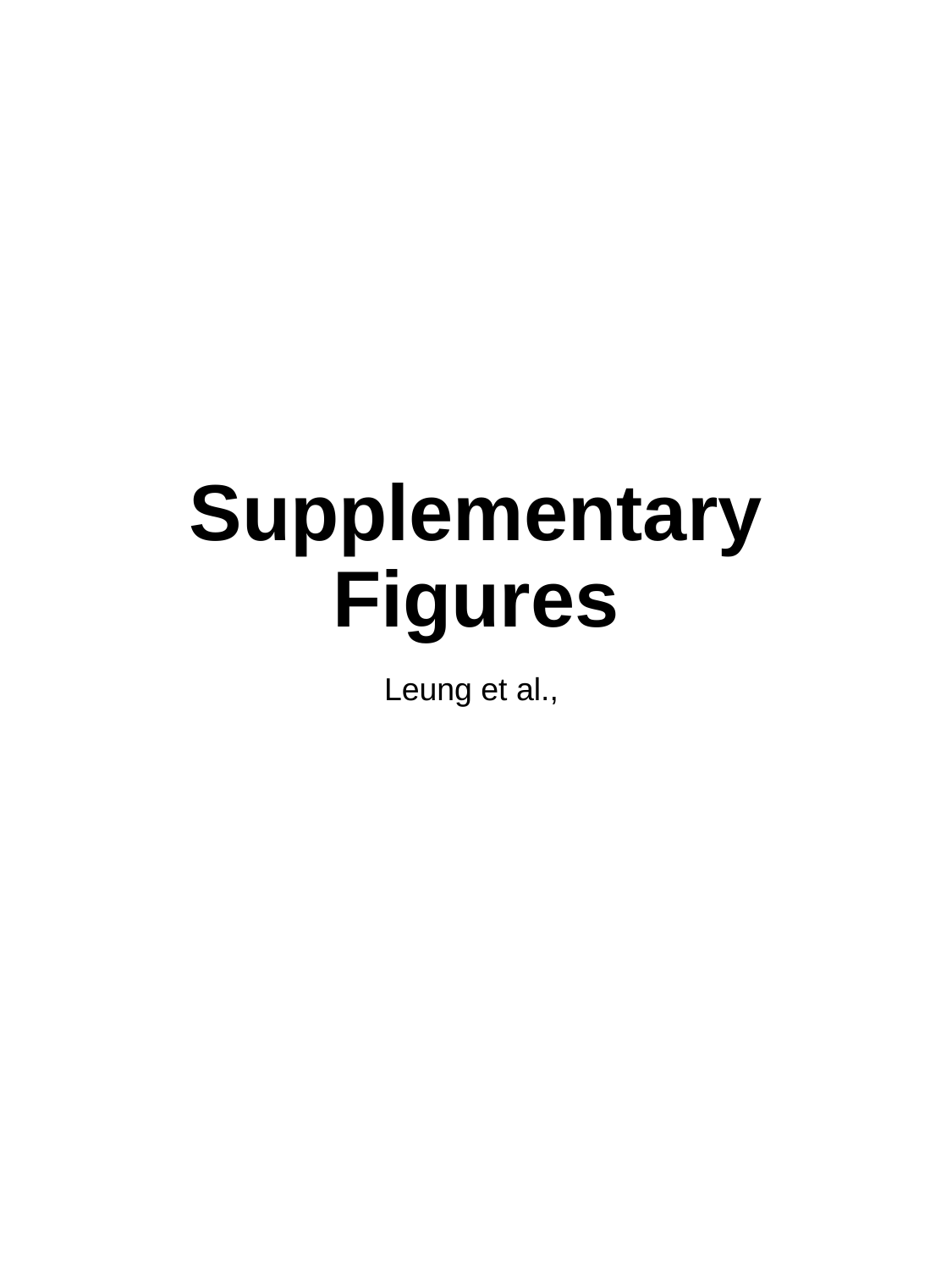

##### Supplementary Figures
Leung et al.,

#### Slide 2
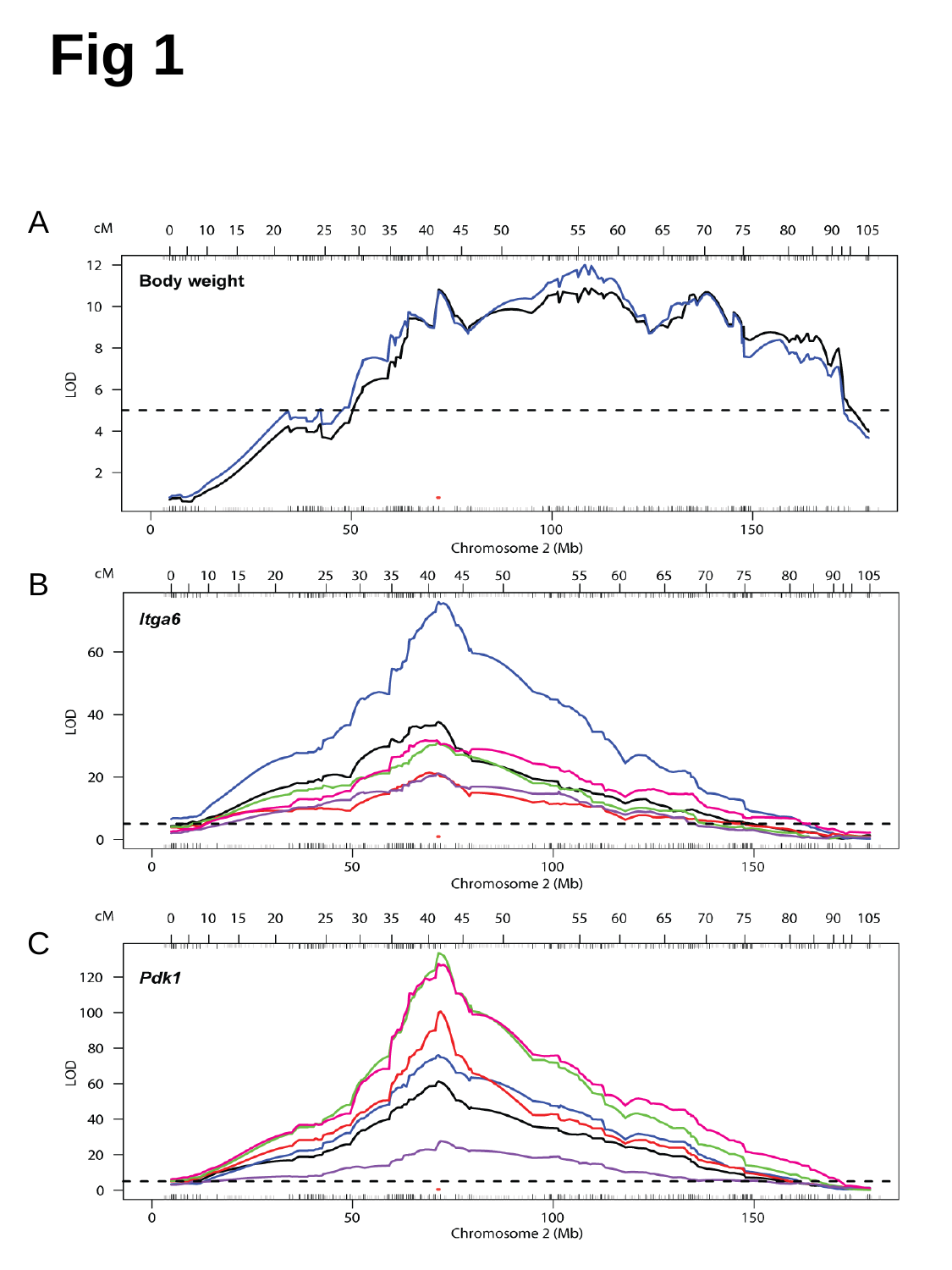

##### Fig 1
A
B
C

#### Slide 3
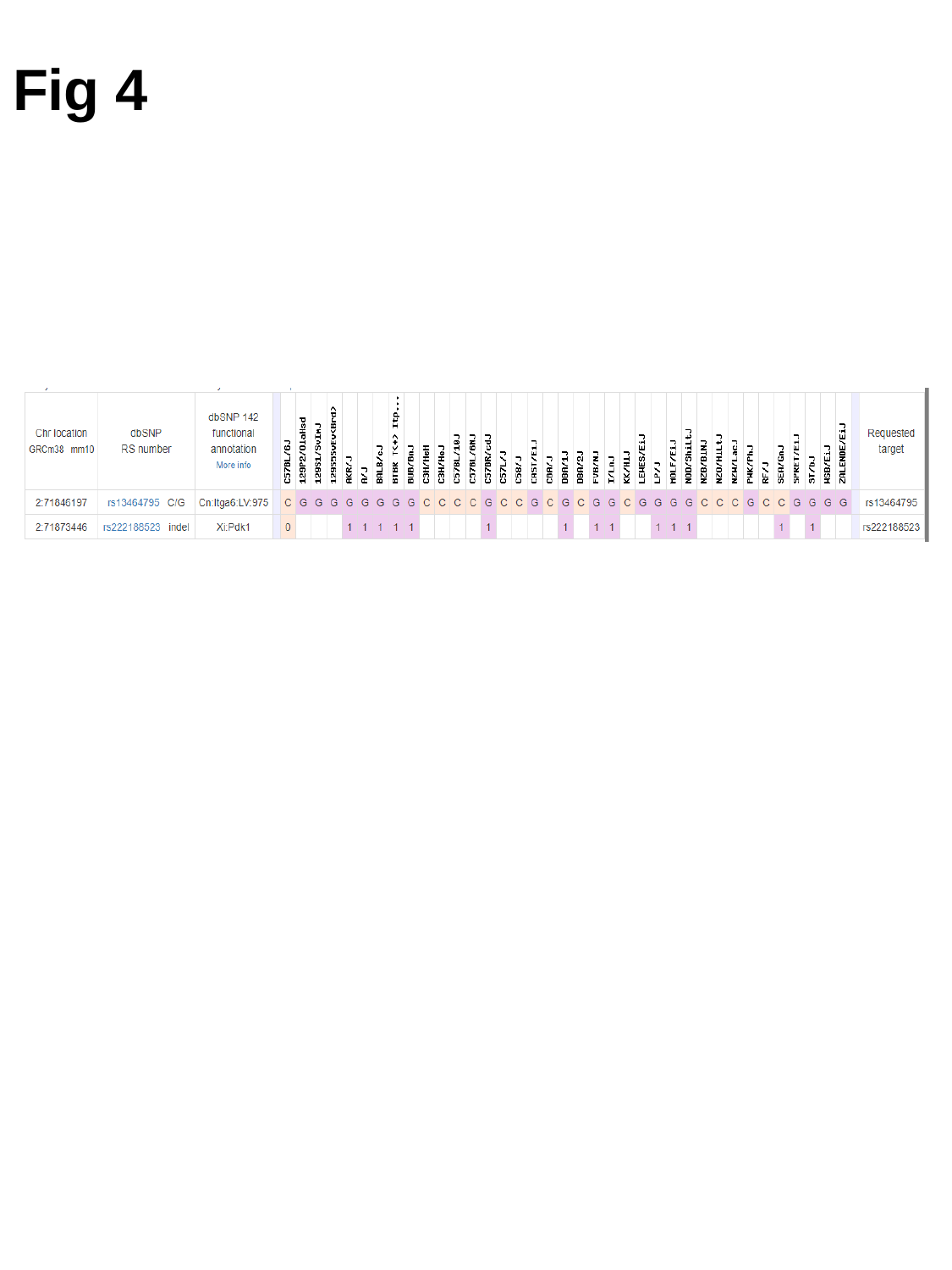

##### Fig 4

#### Slide 4
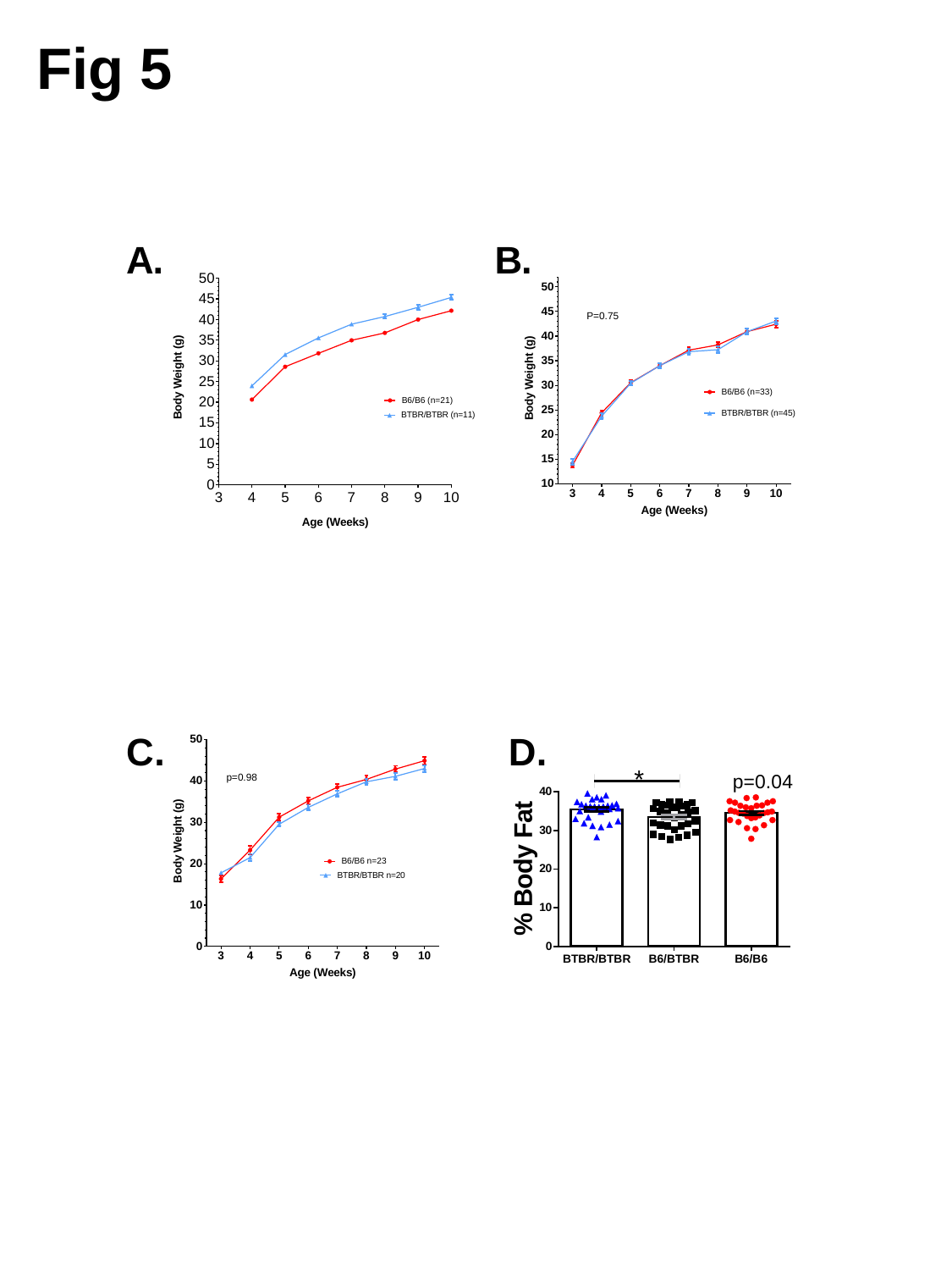

##### Fig 5
